# TR-107, a novel mitochondrial ClpP agonist, induces robust antitumor activity against preclinical models of adrenocortical carcinoma

**DOI:** 10.64898/2026.08.10.743339

**Authors:** George I. Karadimov, Yoo Sun Kim, Haiqing Fu, Sidhant Narula, Fathi Elloumi, Anjali Dhall, Frank Echtenkamp, Luowei Li, Edwin J. Iwanowicz, Lee M. Graves, King Chan, Thorkell Andresson, Robert W. Robey, Yoshimi Greer, Stanley Lipkowitz, Chuong D. Hoang, Jonathan M. Hernandez, Yves Pommier, Mirit I. Aladjem, Urbain Weyemi, Myriem Boufraqech, Suresh M. Kumar, Jaydira del Rivero

## Abstract

Adrenocortical carcinoma (ACC) is a rare and highly aggressive endocrine malignancy originating from the adrenal cortex with limited effective treatment options. The underlying pathophysiology of ACC is uniquely characterized by abnormal steroid production and increased metabolic activity, highlighting the critical role of mitochondria in adrenal steroid hormone biosynthesis and tumor metabolism. In this study, we investigated the therapeutic potential of TR-107, a novel and highly selective small-molecule agonist targeting the mitochondrial protease ClpP. Pharmacologic hyperactivation of ClpP disrupts mitochondrial proteostasis and bioenergetics and has shown promising antitumor activity in various preclinical models. Our results demonstrated that TR-107 induces potent dose-dependent cytotoxic effects at nanomolar concentrations in ACC cell lines NCI-H295R and mACC3 as well as short-term ACC patient-derived organoid (PDO) models, markedly reducing cell viability and confluency *in vitro*. Metabolic analyses revealed that TR-107 significantly impaired oxygen consumption, indicating a disruption of oxidative phosphorylation and substantial attenuation of basal cellular respiration. Mechanistic studies showed dose-dependent increases in reactive oxygen species (ROS) levels and upregulation of proteins involved in mediating the ferroptotic rheostat. Pharmacokinetic assessment uncovered that TR-107 was not a substrate of the ABCB1 (MDR1/P-glycoprotein) efflux transporter, suggesting potential to overcome common multidrug resistance mechanisms. Given the importance of IGF-2 signaling in ACC, we further explored the combinatorial effects of TR-107 with IGF-1 receptor (IGF-1R) inhibitors and discovered that co-treatment produced synergistic reductions in cell viability across NCI-H295R, mACC3, and ACC PDOs. Collectively, these findings support the potential of mitochondrial ClpP hyperactivation as a promising therapeutic strategy for ACC and demonstrate that TR-107 exhibits significant antitumor activity as a monotherapy or in combination with IGF-1R inhibitors. These findings provide a strong rationale for advancing ClpP agonists into clinical development for the management of ACC.

## Introduction

Adrenocortical carcinoma (ACC) is a rare and highly aggressive malignancy originating from the adrenal cortex. The estimated incidence of ACC is approximately 0.7-2 cases per million per year in the general population and approximately 0.2-0.3 per million per year in children and young adults under the age of 20 years (1). ACC exhibits a notable gender predilection, with higher incidence among females than males (55-65% of cases, sex ratio 1.5-2.5:1), and typically follows a bimodal age distribution with peaks in early childhood and between the fourth and fifth decades of life (1–4). Despite its rarity, ACC classically predicts a dismal prognosis, with estimated five-year survival rates varying by ENSAT stage: 65-85% for stage I, 58-68% for stage II, 45-55% for stage III, and 10-20% for stage IV (metastatic) disease (1–4). First-line clinical management of localized ACC is complete surgical resection with negative margins ideally via open adrenalectomy (2,4). However, recurrence is common, with reported rates ranging from approximately 40-80% after radical surgical resection (2,4). The risk of recurrence is modulated by several clinicopathologic factors, including initial disease stage, margin status, Ki67 index, and tumor grade (2,4). Adjuvant mitotane, the only FDA-approved adrenolytic agent for ACC, alone or in combination with etoposide, doxorubicin, and cisplatin (EDP), has remained the cornerstone of systemic therapy for nearly 65 years despite poor tolerability, adrenal cytotoxicity, and persistent uncertainty regarding clinical efficacy (2–6). Mitotane monotherapy demonstrates modest therapeutic benefit in advanced ACC, with an objective response rate of approximately 20-21% and a complete response rate of 3-8% (1,3,4,7). Importantly, mitotane has been shown to induce expression of the ABCG2 (BCRP) drug efflux transporter at the mRNA level (approximately 4-fold induction), with its metabolite o,p’-DDE exhibiting a similar but less potent induction profile *in vitro* (8,9). Furthermore, exposure to mitotane, doxorubicin, and cisplatin has been associated with increased ABCG2 protein expression and drug-resistant phenotypes in ACC cell lines (8,9). Given the poor prognosis associated with ACC and the limited efficacy of available systemic therapies, the development of more effective therapeutic options capable of improving survival, preventing disease recurrence, and combatting metastatic progression remains a critically unmet need.

Metabolic reprogramming has become increasingly recognized as a major oncogenic determinant underpinning aggressive tumor phenotypes including unbridled proliferation, chemotherapeutic resistance, immune evasion, and metastasis (10–15). Tumor-associated metabolic alterations and mitochondrial bioenergetics exert pleiotropic effects, including augmented adenosine triphosphate (ATP) production, preservation of intracellular redox homeostasis, suppression of cell death signaling pathways, and promotion of tumor migration and infiltration in diverse environmental conditions (10–18). Recent work has further identified remarkable metabolic heterogeneity among various tumor types (10,11,14,16,17). While robust glycolytic flux with elevated output of lactate and glycolytic intermediates is nearly ubiquitous among malignancies, reliance on tricarboxylic acid (TCA) cycle intermediates, fatty acid oxidation, amino acid catabolism, glutaminolysis, pentose phosphate pathway (PPP) intermediates, and most notably, oxidative phosphorylation, varies considerably between different tumor types (10–19). In addition, marked tumor metabolic and genotypic plasticity enables cancer cells to exploit multiple metabolic pathways concurrently in distinct contexts (10,11,19,20). For instance, certain tumors exhibit a form of metabolic symbiosis characterized by the concerted and simultaneous utilization of glycolysis and oxidative phosphorylation to sustain growth and survival under hypoxia, enable resistance to antiangiogenic therapies, and adapt to a dynamic tumor microenvironment (20,21). Collectively, this diverse metabolic landscape across tumor types presents unique opportunities for rationally-designed targeted therapeutics that exploit tumor-specific metabolic vulnerabilities.

Oxidative phosphorylation (OXPHOS), the principal mitochondrial pathway responsible for ATP production, has emerged as a critical mediator of tumorigenesis and a particularly compelling therapeutic target (10–12,16,18). Growing evidence from studies among malignancies such as neuroendocrine tumors, gliomas, breast and pancreatic carcinomas, and leukemias has implicated upregulated OXPHOS with multidrug chemotherapeutic resistance, tumor stemness, and proclivity for metastasis (10–12,16,18,20,21). These observations have consequently fueled substantial interest in pharmacologically targeting OXPHOS machinery and disrupting mitochondrial respiratory function to suppress tumor growth and overcome chemotherapeutic resistance (10–12,18,22–26). Consistent with this rationale, several small-molecule inhibitors targeting mitochondrial respiratory complex proteins have displayed notable antitumor efficacy (10–12,23,24,26). These include mitochondrial complex I inhibitors, such as the biguanides metformin and phenformin, quinolone derivative intervenolin, and compounds BAY87-2243, MS-L6, IACS-010759, and ASP4132, all of which demonstrated potent activity across *in vitro* and *in vivo* tumor models (10–12,23–26). However, clinical translation of these agents has been hampered by significant toxicity. BAY 87-2243 entered a phase I trial (NCT01897116) that was terminated early due to dose-independent emesis (27,28). IACS-010759 was evaluated in two phase I trials in relapsed/refractory AML (NCT02882321) and advanced solid tumors (NCT03291938), where it demonstrated a narrow therapeutic index with dose-limiting elevated blood lactate and neurotoxicity; no recommended phase 2 dose was established, and both trials were discontinued (28). ASP4132 was evaluated in a phase I dose-escalation study (NCT02383368) in 39 patients with advanced solid tumors, where dose-limiting toxicities including fatigue, mental status changes, lactic acidosis, and posterior reversible encephalopathy syndrome prohibited dose escalation beyond 5 mg, and only stable disease was observed in 20.5% of patients (29). Ultimately, these outcomes highlight that novel approaches to target mitochondrial respiratory machinery in OXPHOS-dependent tumors are urgently needed.

Caseinolytic protease P (ClpP) is a barrel-shaped tetradecameric serine protease located in the mitochondrial matrix of a diverse range of eukaryotic organisms (11,12,30,31). While ClpP possesses intrinsic ATP-independent peptidase activity, its full proteolytic function requires assembly into a stable ClpXP complex with its hexameric chaperone ClpX, an ATP-dependent unfoldase and polypeptide translocase itself a member of the AAA+ protein superfamily (11,12,30–33). Analyses of the ClpXP crystal structure indicate that ClpX engages with certain hydrophobic pockets on ClpP to induce conformational changes that remodel electrostatic interactions and facilitate opening of the ClpP axial entrance pores (11,12,30–34). In this activated state, ClpX mediates the selective recognition, unfolding, and translocation of polypeptide substrates threaded into the proteolytic chamber of ClpP for consequent degradation (11,12,30–34). The ClpXP complex is widely recognized as a canonical mediator of the mitochondrial unfolded protein response, where it regulates proteostasis through the recognition and degradation of misfolded or damaged mitochondrial proteins (11,12,30–35). However, the multiplicity of ClpXP substrates, including respiratory chain subunits, TCA cycle proteins, and mitochondrial DNA (mtDNA) transcription and translation factors, reflects a multifaceted yet central role in mitochondrial homeostasis (11,12,34).

ClpP constitutes a unique target for cancer therapy, as both inhibition and hyperactivation of its proteolytic activity have demonstrated antitumor efficacy (33,36). Pharmacologic inhibition of ClpP through small-molecules primarily targeting the canonical serine-histidine-aspartate catalytic triad results in the aggregation of damaged and misfolded respiratory chain proteins that disrupt OXPHOS, promote the production of mitochondrial reactive oxygen species, and initiate apoptosis (30,36,37). Conversely, ClpP hyperactivation induces uncontrolled proteolysis of its substrates, thereby leading to impaired OXPHOS, attenuated TCA cycle metabolism, and subsequent cell death (11,22,30,36,37). In contrast to ClpP inhibitors, ClpP agonists displace ClpX at its interface with ClpP hydrophobic pockets and enlarge the axial entrance pore to configure a ClpX-independent hyperactive state (11,12,22,30,33,36,37). Acyldepsipeptides (ADEPs) and subsequent analogs were the first known ClpP agonists and demonstrated substantial antibiotic activity against multidrug-resistant bacteria (11,12,38). More recently, the first-in-class small-molecule imipridones ONC201 and ONC212, originally identified through screens for compounds capable of stimulating TNF-related apoptosis-inducing ligand (TRAIL), have emerged as potent antitumor agents and highly selective ClpP agonists (11,12,22,30,37,39–42). ONC201, in particular, has exhibited antitumor activity *in vitro* and *in vivo* across a broad spectrum of aggressive malignancies, including acute myeloid leukemia, breast cancer, and gliomas, with minimal sensitivity in non-malignant cells (11,12,22,30,33,37,39–43). ONC201 (dordaviprone) received FDA accelerated approval on August 6, 2025, for adult and pediatric patients ≥1 year of age with H3 K27M-mutant diffuse glioma with progressive disease following prior therapy, based on an overall response rate of 22% (BICR, RANO 2.0) with a median duration of response of 10.3 months, representing the first FDA-approved systemic therapy for this indication (11,43–45). These findings have also led to the initiation of the ACTION trial, an international randomized, double-blind, placebo-controlled phase III clinical trial of dordaviprone in patients with newly diagnosed H3 K27M-mutant diffuse glioma (NCT05580562) (46).

TR-107 (Fig. 1A) is a next-generation ClpP agonist with significantly improved potency, selectivity, binding affinity, oral bioavailability, and pharmacokinetic stability compared to imipridone forebears ONC201, ONC206, and ONC212 (11,12). Mechanistic investigation has shown that TR-107 hyperactivates ClpP, resulting in depletion of mitochondrial respiratory proteins, disruption of OXPHOS, and profound mitochondrial dysfunction (11,12). While the antitumor effect of TR-107 was initially characterized in triple-negative breast cancer cell lines and xenograft models, subsequent studies in colorectal cancer and glioblastoma models have further validated potent nanomolar activity in both monotherapy and combination therapy (11,12,47). Herein, we investigate the efficacy of TR-107 against ACC using NCI-H295R, mACC3, and novel short-term ACC patient-derived organoid (NCI-ACC PDO) models. We first establish the dose-dependent cytotoxicity of TR-107 in ACC. Next, we examine the mechanism of action, including profiling and assessing mitochondrial disruption and metabolic impairment, followed by characterizing a predominant mode of cell death in both NCI-H295R and NCI-ACC PDOs. Thereafter, we determine whether TR-107 serves as a substrate of the ATP-binding cassette transporter B1 (ABCB1/MDR1/P-glycoprotein) drug efflux transporter. Finally, we investigate the combinatorial potential of TR-107 with IGF-1R inhibition to assess synergistic antitumor efficacy and establish promising therapeutic modalities adaptable to the clinical management of ACC.

**Figure 1.**
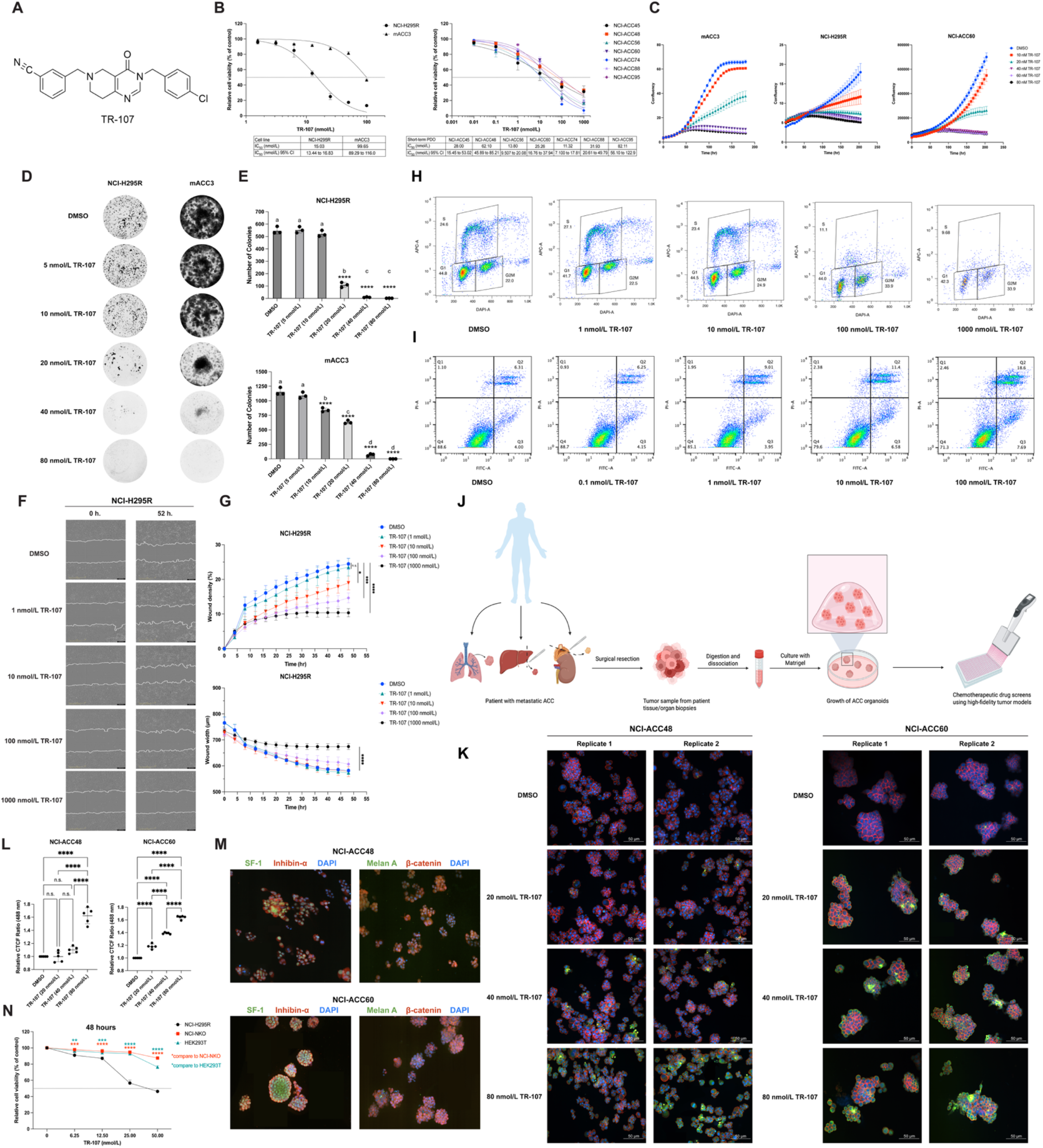
TR-107 exhibits potent cytotoxicity against preclinical models of adrenocortical carcinoma. **A,** Chemical structure of TR-107 designed using ChemDraw (Revvity). **B,** Dose-response curve of TR-107 following 72-hour treatment across NCI-H295R, mACC3, and multiple NCI-ACC PDOs with corresponding IC_50_ values in nmol/L. Cell viability was normalized to vehicle-treated controls and expressed as a percentage of vehicle controls. **C,** Longitudinal cell proliferation of NCI-H295R, mACC3, and NCI-ACC60 following treatment with the indicated concentrations of TR-107. Data reported as mean ± SEM, *n* = 3 technical replicates. **D** and **E,** Representative crystal violet staining (**D**) and quantification of NCI-H295R and mACC3 colony formation (**E**) following TR-107 treatment. **F** and **G,** Representative images from NCI-H295R scratch wound assays (**F**) and quantification of wound density and width (**G**) following TR-107 treatment. **H** and **I,** Flow cytometric analyses of EdU incorporation (**H**) and Annexin V/propidium iodide (PI) staining (**I**) in NCI-H295R cells following TR-107 treatment. **J,** Schematic overview of short-term NCI-ACC PDO generation created using BioRender.com. **K** and **L,** Representative fluorescent images of NCI-ACC48 and NCI-ACC60 PDOs following TR-107 treatment (**K**) and quantification of green fluorescence (**L**). Green, Image IT-DEAD reagent; red, phalloidin; blue, DAPI. Quantification was performed from five independent fluorescence measurements per condition. For each measurement, fluorescence intensity values from two biological replicates were averaged and reported as individual data points. **M,** Representative immunofluorescence staining demonstrating expression of characteristic ACC markers in both NCI-ACC48 and NCI-ACC60 PDOs. **N,** Analysis of TR-107 selectivity using NCI-H295R, HEK293T, and normal kidney PDO models. Following 48-hour TR-107 treatment at the indicated concentrations, cell viability was normalized to vehicle-treated controls and expressed as a percentage of vehicle controls. Data reported as mean ± standard deviation, *n* = 3 technical replicates unless otherwise specified. n.s., not significant; **, *P* < 0.01; ***, *P* < 0.001; ****, *P* < 0.0001. ACC, adrenocortical carcinoma; PDO, patient-derived organoid.

## Materials and Methods

### Cell lines

Human ACC cell line NCI-H295R (CRL-2128) was obtained from American Type Culture Collection (ATCC) and cultured in 1:1 Dulbecco’s Modified Eagle Medium/Nutrient Mixture F-12 (Gibco, #11320033) containing 2.5% Nu-Serum (Corning, #355100), 1% ITS + Premix Universal Culture Supplement (Corning, #354352), and 1% penicillin-streptomycin. Murine ACC cell line mACC3 was obtained from the cell line repository at the Developmental Therapeutics Branch of the National Cancer Institute (NCI). This cell line was derived from a genetically engineered murine model harboring loss-of-function (LOF) mutations in *Trp53* and gain-of-function (GOF) mutations in *Ctnnb1* (β-catenin), which develops ACC as previously described (48). mACC3 was cultured in 1:1 Dulbecco’s Modified Eagle Medium/Nutrient Mixture F-12 (Gibco, #11320033) containing 2.5% Nu-Serum (Corning, #355100), 1% ITS + Premix Universal Culture Supplement (Corning, #354352), and 1% penicillin-streptomycin.

Human embryonic kidney cell line HEK293T (CRL-3216) was obtained from ATCC and cultured in Dulbecco’s Modified Eagle Medium (Gibco, #11995065) containing 1% penicillin-streptomycin. All cell lines were maintained at 37°C in a humidified incubator with 5% CO_2_, regularly tested for mycoplasma contamination using the MycoAlert Mycoplasma Detection Kit (Lonza, #LT07-318), and authenticated by short tandem repeat (STR) profiling (Laragen Inc.)

### Patient tumor specimens

Fresh surgically resected patient tumors were obtained from patients treated at the National Institutes of Health (NIH) Clinical Center and were used for patient-derived organoid (PDO) generation and experimentation. Tumor specimens were obtained from either primary adrenal tumor or metastatic lesions at the time of clinically indicated surgical resection. All patients were provided written informed consent under Institutional Review Board (IRB)-approved protocols NCT05237934 and NCT06041516. Clinical and molecular characteristics of the patient cohort are summarized in Table 1. This study complied with all applicable institutional guidelines and ethical regulations.

**Table 1.**
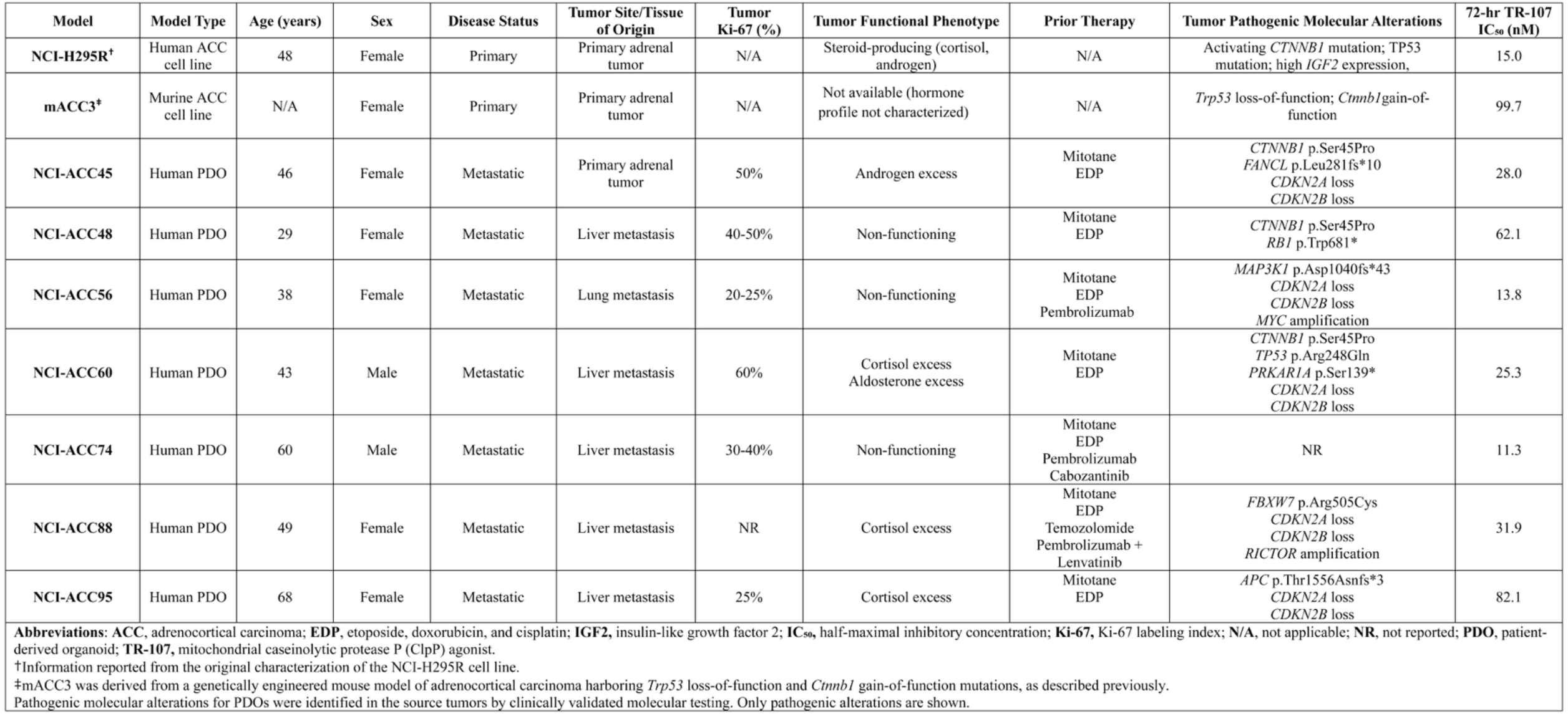
Clinical, pathological, and molecular characteristics of experimental ACC models.

### Short-term patient-derived organoids (PDOs)

Short-term NCI-ACC PDOs were generated using fresh surgically resected ACC tumor specimens obtained under protocols NCT05237934 and NCT06041516. Tumors were mechanically and enzymatically dissociated into single cells as previously described (49,50). Single tumor cells were resuspended in Growth-Factor Reduced Matrigel (Corning, #354230) and plated in gamma irradiation-sterilized tissue culture-treated plates. Tumor-Matrigel suspensions were maintained at 37°C in a humidified incubator with 5% CO_2_ for 20 minutes to allow solidification. Following polymerization, tumor-Matrigel dome suspensions were overlaid with Minimum Basal Media (MBM) composed of 1:1 Dulbecco’s Modified Eagle Medium/Nutrient Mixture F-12 (Gibco, #11320033) containing 1x N2 supplement (Gibco, #17502048), 1x B27 supplement (Gibco, #17504044), 50 ng/mL recombinant human EGF (STEMCELL Technologies, #78006), 20 ng/mL recombinant human bFGF (STEMCELL Technologies, #78003), 100 ng/mL recombinant human IGF-2 (STEMCELL Technologies, #78023), 10 μM Y-27632 (STEMCELL Technologies, #72304), and 100 U/mL penicillin-streptomycin. PDOs were passaged every 2 to 4 weeks through gentle mechanical dissociation using Dispase in DMEM/F12 (STEMCELL Technologies, #07923) followed by enzymatic dissociation into single cells using 0.05% Trypsin-EDTA (Gibco, #25-300-054). NCI-ACC PDO cells were subsequently re-embedded in Matrigel with fresh MBM for continued propagation or experimental use.

### *In vitro* cell line and PDO cytotoxicity assays

Single cell suspensions from human or murine ACC cells, human embryonic kidney cells, or human adrenal cells were seeded into opaque 384-well plates in triplicate at a density of 2×10^3^cells per well in 30 µL of respective culture medium. Single cells from NCI-ACC PDOs were seeded embedded in 10 µL of Growth-Factor Reduced Matrigel in triplicate, incubated at 37°C until solidification, and finally overlaid with 20 µL of fresh MBM medium. Both cells and NCI-ACC PDOs were permitted to adhere for 48 hours at 37°C in a humidified incubator with 5% CO_2_. Following cell adherence and equilibration, cells or NCI-ACC PDOs were treated with 30 µL of fresh medium containing vehicle control (dimethyl sulfoxide [DMSO]) or variable concentrations and combinations of investigational compounds for 48 or 72 hours. At the conclusion of the indicated treatment periods, relative cell viability was assessed using the CellTiter-Glo Luminescent Cell Viability Assay (Promega, #G7573). Briefly, 20 µL of CellTiter-Glo reagent was added to each well and luminescence was quantified with the SpectraMax i3x Multi-Mode Microplate Reader (Molecular Devices) according to the manufacturer’s instructions. Synergistic cytotoxic drug interactions in cell lines and NCI-ACC PDOs were evaluated and analyzed using the standalone Windows software Combenefit (http://sourceforge.net/projects/combenefit/) (51). Synergy was calculated using the Highest Single Agent (HSA) method, a mathematical model that compares the observed combinatorial effect with that of the most effective single agent at the corresponding concentration. For additional investigation into synergistic drug interactions, NCI-ACC48 and NCI-ACC60 PDOs were treated with variable concentrations of respective compounds for the indicated 7- or 14-day treatment period. NCI-ACC PDOs were subsequently visualized using the Olympus IX73 Standard Inverted Microscope (Evident Scientific) and captured with the cellSens Imaging Software (Evident Scientific).

For rescue experiments using ferroptosis inhibitors, ABCB1/ABCG2 inhibitors, and antioxidant compounds, NCI-H295R and NCI-ACC PDOs were seeded in 384-well plates as described above and maintained at 37°C in a humidified incubator with 5% CO_2_ for 48 hours. Thereafter, NCI-H295R or NCI-ACC PDOs were pretreated for 6 hours with fresh medium containing respective inhibitory or antioxidant compounds at the indicated concentrations prior to the addition of fresh medium comprising variable concentrations of TR-107. Relative cell viability was assessed 48 hours after TR-107 treatment using the CellTiter-Glo Luminescent Cell Viability Assay (Promega, #G7573) and quantified using the SpectraMax i3x Multi-Mode Microplate Reader (Molecular Devices) as described above.

### shRNA-mediated gene silencing and *in vitro* growth assays

*IGF2* shRNA lentiviral particles (Santa Cruz Biotechnology, sc-39576-V), *IGF1R* shRNA lentiviral plasmid pLV[shRNA]-Puro-U6-hIGFR#1 (VB260326-1546dah) (VectorBuilder Inc.), and control shRNA lentiviral particles-A (Santa Cruz Biotechnology, sc-108080) were purchased and used according to the manufacturer’s instructions. NCI-H295R cells were transduced with *IGF2* shRNA lentiviral particles, *IGF1R* shRNA lentiviral plasmid, or control shRNA lentiviral particles at approximately 60% to 80% confluence in complete growth medium containing FBS for 24 hours at 37°C. Transduced cells were selected using 1 µg/mL puromycin, and stable puromycin-resistant cells were expanded and maintained for further experimentation. For growth assays, NCI-H295R, mACC3, NCI-H295R *IGF2* shRNA, NCI-H295R control shRNA control and PDOs NCI-ACC48 and NCI-ACC60 were seeded in triplicate technical replicates as described above in a clear-bottom 96-well plate at a density of 2×10^3^ cells per well. Following a 24-hour incubation period at 37°C with 5% CO_2_, cells were overlaid with fresh medium containing vehicle control DMSO or variable concentrations and combinations of investigational compounds. Live cell proliferation was monitored using the Incucyte S3 Live-Cell Analysis Instrument (Sartorius, Ann Arbor, MI). Phase contrast images were acquired every 6 hours at 10x or 4x magnification, and confluency (μm²/image) was recorded and quantified over time.

For clonogenic assays, NCI-H295R and mACC3 cell lines were seeded in triplicate technical replicates at a density of 1×10^3^cells per well in 12-well plates pre-coated with 0.1% gelatin in 1x PBS. After 24 hours of cell adherence, NCI-H295R and mACC3 cells were continuously treated with indicated compounds for 14 or 10 days, respectively. Colonies were fixed with methanol and stained with 0.1% crystal violet in methanol for 30 minutes. Staining solution was aspirated and the colonies were imaged using ChemiDoc MP Imaging System (Bio-Rad) and counted using ImageJ (https://github.com/imagej/ImageJ) and OpenCFU (https://opencfu.sourceforge.net/) (52,53). For scratch wound assays, Incucyte ImageLock 96- well plates (Sartorius, Ann Arbor, MI) were precoated with 50 µL of poly-D-lysine and NCI- H295R cells were seeded in triplicate biological replicates at 3×10^4^ cells per well and maintained at 37°C in a humified incubator with 5% CO_2_ overnight. At 95% to 100% confluence, uniform scratch wounds were created using the Incucyte 96-Well Woundmaker Tool (Sartorius, Ann Arbor, MI). Following wound introduction to the adherent cell monolayer, 2 mg/mL mitomycin C was added for 2 hours to inhibit proliferation before replacement with fresh medium containing vehicle control DMSO or TR-107 at variable concentrations. Phase contrast images were acquired at 10x magnification every 6 hours using the Incucyte S3 Live-Cell Analysis Instrument (Sartorius, Ann Arbor, MI) for a period of 48 hours.

### Flow cytometry and apoptosis assays

For EdU incorporation studies, NCI-H295R cells were processed using the Click-iT Plus EdU Alexa Fluor 647 Flow Cytometry Assay Kit (Invitrogen, #C10634) according to the manufacturer’s instructions. Briefly, 2×10^5^ NCI-H295R cells were plated in tissue-culture 6-well plates and allowed to adhere overnight. Cells were then treated with fresh medium containing vehicle control DMSO or variable concentrations of TR-107 for 48 hours. Upon completion of the treatment period, cells were labeled with 10 µM Click-iT EdU and incubated for 1 hour at 37°C with 5% CO_2_. Cells were fixed with 4% paraformaldehyde (PFA) in PBS and permeabilized with 3% bovine serum albumin (BSA) and 0.5% Triton X-100. Click-iT Plus reaction cocktail was subsequently added and cells were incubated for 30 minutes at room temperature protected from light. Cells were then stained with 2 µg/mL of DAPI and detected using an LSR Fortessa cytometer (BD Biosciences) and analyzed using FlowJo software version 10.10.0. For drug-induced apoptosis studies, NCI-H295R cells were processed as per the manufacturer’s instructions using the FITC Annexin V Apoptosis Detection Kit with PI (BioLegend, #640914). Briefly, 2×10^5^ NCI-H295R cells were plated in 6-well tissue culture plates, allowed to adhere over 48 hours, and thereafter overlaid with fresh medium containing vehicle control DMSO or variable concentrations of TR-107 for 48 hours. Cells were harvested, washed with ice-cold PBS containing 1% BSA, and resuspended in Annexin V binding buffer. Following incubation with FITC-conjugated Annexin V for 15 minutes at room temperature protected from light, propidium iodide (PI) was added immediately prior to data acquisition. Data were acquired using an LSR Fortessa cytometer (BD Biosciences) and analyzed using FlowJo software version 10.10.0.

For the Image-IT DEAD Green viability fluorescent staining, NCI-ACC48 and NCI- ACC60 PDOs were cultured in Lab-Tek 8-well chamber glass slides (Thermo Scientific, # 154941PK) in biological replicates at a density of 2×10^3^ cells per well as described above. Upon reaching approximately 80% confluence, NCI-ACC48 and NCI-ACC60 PDO were overlaid with fresh medium containing vehicle control DMSO or variable concentrations and combinations of TR-107 or ceritinib for 48 hours. Following completion of the treatment period, 100 nM of the Image-IT DEAD Green reagent (Thermo Scientific, #I10291) was added to the NCI-ACCC48 and NCI-ACC60 PDO cultures and incubated for 4 hours at 37°C with 5% CO_2_. Drug-treated PDOs were fixed with 4% PFA in PBS with 1% BSA followed by permeabilization with 0.3% Triton X-100, 0.05% Tween 20, and 1% BSA in ice-cold PBS. Fixed and permeabilized PDOs were stained with 0.1 ug/mL TRICTC phalloidin and 2 µg/mL of DAPI solution. Fluorescent images were acquired using confocal microscopy (Nikon CSU-W1 SoRa). Fluorescence was quantified using ImageJ (https://github.com/imagej/ImageJ) by calculating the corrected total cell fluorescence (CTCF), defined as integrated density − (cell area × mean background fluorescence).

### Immunofluorescence staining

NCI-ACC48 and NCI-ACC60 PDOs were cultured in Lab-Tek 8-well chamber glass slides (Thermo Scientific, # 154941PK) at a density of 2×10^3^ cells per well as described above. Upon reaching approximately 80% confluence, NCI-ACC48 and NCI-ACC60 PDOs were fixed with 4% PFA in ice-cold 1x PBS. Fixed samples were washed with ice-cold PBS and thereafter permeabilized and blocked for 2 hours with 5% goat serum (Gibco, #16210064) and 0.5% Triton X-100 in PBS at room temperature. Fixed and permeabilized PDOs were then incubated overnight at 4°C with following primary antibodies at a 1:100 dilution: SF-1 (A-1) (Santa Cruz Biotechnology, #sc-393592, mouse, monoclonal), Melan A (A103) (Santa Cruz Biotechnology, #sc-20032, mouse, monoclonal), β-catenin (D10A8) (Cell Signaling Technology, #8480S, rabbit, monoclonal), Inhibin-α (Bioss, #bs-1032R, rabbit, polyclonal), and GPX4 (Cell Signaling Technology, #52455, rabbit, polyclonal). Following primary antibody incubation, PDOs were washed with ice-cold PBS and incubated with fluorophore-conjugated secondary antibodies at a 1:500 dilution for 2 hours at room temperature protected from light. Fluorophore-conjugated secondary antibodies included Alexa Fluor 488 goat anti-mouse IgG (H+L) (Invitrogen, #A- 11001, polyclonal), Alexa Fluor 568 goat anti-rabbit IgG (H+L) (Invitrogen, #A-11036, polyclonal), Alexa Fluor 488 (Invitrogen, #A12379), and rhodamine phalloidin (Invitrogen, #R415). PDOs were subsequently washed with ice-cold PBS and stained with 2 µg/mL of DAPI. Fluorescent images were acquired using confocal microscopy (Nikon CSU-W1 SoRa).

### Mitochondrial respiration

For analysis of mitochondrial stress, NCI-H295R cells were seeded at the empirically optimized cell density of 3×10^4^ cells per well in poly-L lysine treated Seahorse XFe96/XF Pro FluxPak Cell Culture Microplates (Agilent Technologies, #103792-100) and incubated at 37°C in a humidified incubator with 5% CO_2_ for 24 hours. After incubation, cells were overlaid with fresh medium containing vehicle control DMSO or variable concentrations of TR-107 and treated for 48 hours. Mitochondrial respiration was then assessed using the Seahorse XF Cell Mito Stress Test Kit (Agilent Technologies, #103015-100). Briefly, the medium was replaced with XF DMEM pH 7.4 supplemented with fresh 10 mM glucose, 1 mM sodium pyruvate, and 2 mM glutamine. Oxygen consumption (OCR) and extracellular acidification rate (ECAR) were obtained using the Seahorse XFe96/XF96 Extracellular Flux Analyzer (Agilent Technologies). Mitochondrial stress test assays were conducted with the following final concentrations of inhibitors: 1 µM oligomycin, 1 µM FCCP, and 0.5 µM rotenone/antimycin A. Samples were normalized by CyQUANT (ThermoFisher Scientific, #C7026). Analysis was conducted on Wave 2.6.3 (Agilent) and data presented are the result of at least four wells per condition and reported as the mean ± the standard deviation of rates.

### Enzyme-linked immunosorbent assays (ELISAs)

Analyses of testosterone, cortisol, and IGF-2 secretion were performed using the Human Testosterone (Competitive EIA) ELISA Kit (LS Bio, #LS-F10017-1), Human Cortisol (Competitive EIA) ELISA Kit (LS Bio, #LS-F10024-1), and the Human IGF-II/IGF2 Quantikine ELISA Kit (R&D Systems, #DG200), respectively, as per the manufacturer’s instructions. For testosterone and cortisol ELISAs, NCI-H295R and NCI-ACC60 PDOs were seeded in biological quadruplicates in tissue-culture 6-well plates as described above. Cultures were maintained at 37°C in a humidified incubator with 5% CO_2_ until they achieved approximately 80-85% confluence. The medium was then replaced with fresh medium containing either vehicle control DMSO or variable concentrations of TR-107, and cultures were incubated for an additional 48 hours. Following completion of the treatment period, conditioned medium was collected and centrifuged at 462×g for 5 minutes to remove cellular debris. Clarified supernatant was then used for subsequent ELISA analyses. Absorbance values were measured at 450 nm with 540 nm background correction using the SpectraMax i3x Multi-Mode Microplate Reader (Molecular Devices). Testosterone and cortisol concentrations and ratios were calculated according to the manufacturer’s instructions. For IGF-2 ELISAs, NCI-H295R, NCI-ACC48, and NCI-ACC60 PDOs were seeded as described above in quadruplicate biological replicates across two independent experiments. Cells were maintained at 37°C in a humidified incubator with 5% CO_2_ until they achieved approximately 80-85% confluence. The culture medium was then replaced with fresh growth medium without hormone supplementation, and conditioned medium was collected 24 hours later. Following centrifugation at 462×g for 5 minutes, the clarified supernatant was analyzed by ELISA. Absorbance values were measured at 450 nm with 540 nm background correction using the SpectraMax i3x Multi-Mode Microplate Reader (Molecular Devices). Basal IGF-2 concentrations were calculated according to the manufacturer’s instructions.

### Targeted metabolomics

Targeted metabolomics was performed by organic solvent extract from NCI-H295R and NCI-ACC60 cells following 24-hour drug treatment using stable isotope dilution LC-MS/MS based targeted multiple reaction monitoring (MRM) assays. Metabolite extraction was performed by adding 500 µL chilled 80% methanol-water solution to cell pellets. Samples were vortexed vigorously for 30 seconds and centrifuged at 14,000 × g for 10 min. Following centrifugation, 120 µL of supernatant was transferred to an autosampler vial. Samples were dried in a SpeedVac vacuum concentrator (ThermoFisher Scientific) and then reconstituted in 40 µL of water. Five µL of sample was injected for reversed-phase ion-pairing LC-MS^2^ analysis. The pellets generated during the metabolite extraction were solubilized in EasyPep lysis buffer (ThermoFisher Scientific) supplemented with DNase and then used for protein estimation and data normalization. Selected metabolites in glycolysis, pentose phosphate, TCA cycle, oxidative phosphorylation and redox pathways were measured and quantified. LC-MS analysis was performed on Thermo TSQ Quantiva triple quadrupole mass spectrometers (Thermo Scientific, San Jose, CA) coupled to a NexeraXR LC system (Shimadzu Scientific Instruments, Columbia, MD). Both the HPLC and mass spectrometer were controlled by Xcalibur software (Thermo Scientific). Quantitation of all metabolites was carried out using Xcalibur Quan Browser (Thermo Scientific). Calibration curves for each metabolite were constructed by plotting the ratio of the reference compound over the spiked in isotopic standard peak area ratios obtained from the calibration standards curve and fitting the data using linear regression with 1/*X* weighting. The metabolite concentrations in samples were then interpolated using the linear function obtained from the calibration curve.

### Oxidative stress and lipid peroxidation assays

NCI-H295R cells and NCI-ACC60 PDOs were dissociated into single-cell suspensions and seeded at a density of 2×10^5^ cells per well in tissue-culture 6-well plates in triplicate biological replicates. Cultures were maintained at 37°C with 5% CO_2_ in a humidified incubator until adherent monolayers were established. Upon reaching approximately 85% confluence, cells and PDOs were overlaid with fresh medium containing vehicle control DMSO or variable concentrations of TR-107 for 48 hours. Cellular ROS was then analyzed and quantified using the DCFDA/H2DCFDA Cellular ROS Assay Kit (Abcam, #ab113851) according to the manufacturer’s instructions. Briefly, DCFDA was added to drug-treated cells and PDOs in a final concentration of 20 µM exactly 30 minutes prior to completion of drug treatment. Cultures were then incubated for an additional 30 minutes at 37°C with 5% CO_2_ in a humidified incubator. Fluorescence intensity was immediately obtained and quantified using the SpectraMax i3x Multi-Mode Microplate Reader (Molecular Devices) with ex/em = 485/535 nm. Lipid peroxidation was quantified by measuring malondialdehyde (MDA) using the Lipid Peroxidation (MDA) Assay Kit (RayBiotech, #MA-MDA-2) in accordance with the manufacturer’s instructions. In short, cells and PDOs were seeded as described above, cultured to approximately 85% confluence, and overlaid with fresh medium containing vehicle control DMSO or variable concentrations of TR-107 for 48 hours. Following completion of the treatment period, cells and PDOs were washed twice with ice-cold 1x PBS, pelleted by centrifugation at 1,000×g for 10 minutes, and lysed in 1x lysis buffer supplemented with 100x BHT solution. Cell lysates were then incubated on a rocker at 4°C for 30 minutes, followed by centrifugation at 14,000×g for 10 minutes at 4°C. Clarified protein lysate was collected for subsequent lipid peroxidation analysis. Thereafter, 100 µL of MDA standard or clarified protein lysate was transferred to a microcentrifuge tube and mixed with 100 µL of SDS solution. Subsequently, 250 µL of prepared thiobarbituric acid (TBA) reagent was added to each standard and sample, mixed thoroughly, and incubated at 95°C for 60 minutes. Following incubation, samples were cooled on ice and centrifuged at 1,600×g for 10 minutes. The resulting supernatant was transferred to a clear-bottom 96-well plate and absorbance was quantified at 532 nm immediately using the SpectraMax i3x Multi-Mode Microplate Reader (Molecular Devices). MDA concentrations were determined according to the manufacturer’s instructions.

NCI-H295R mito-roGFP2 cells were generated by retroviral transduction using replication-incompetent HEK293FT-derived retrovirus encoding mito-roGFP2-Orp1 in accordance with previously published protocols (54,55). NCI-H295R cells were seeded in tissue-culture 6-well plates, allowed to adhere overnight, and transduced with mito-roGFP2 retrovirus in the presence of 10 µg/mL of polybrene. After 24 hours, viral supernatant was replaced with fresh growth media, and cells were cultured for an additional 48 hours prior to puromycin selection. NCI-H295R mito-roGFP2 cells were selected using 1 µg/mL of puromycin, and stable puromycin-resistant cells were isolated and expanded. NCI-H295R mito-roGFP2 cells were subsequently seeded in triplicate biological replicates at 2×10^5^ cells per well in tissue-culture 6- well plates and treated with variable concentrations of TR-107 or indicated compounds. Time-lapse fluorescent imaging was initiated immediately following compound addition and continued overnight using confocal microscopy (Nikon CSU-W1 SoRa). Fluorescence was quantified with ImageJ (https://github.com/imagej/ImageJ) by calculating the corrected total cell fluorescence (CTCF), defined as integrated density − (cell area × mean background fluorescence).

### Quantitative reverse transcription polymerase chain reaction (RT-qPCR)

NCI-H295R and NCI-ACC60 PDO cells were dissociated into single-cell suspensions and seeded at a density of 2×10^5^ cells per well in tissue-culture 6-well plates in triplicate biological replicates. Cultures were maintained at 37°C with 5% CO_2_ in a humidified incubator until adherent monolayers were established. Upon reaching approximately 85% confluence, cells and PDOs were overlaid with fresh medium containing vehicle control DMSO or variable concentrations of TR-107 for 24 hours. Following completion of the treatment period, RNA isolation was performed using the RNeasy Mini Kit (QIAGEN, #74104) according to the manufacturer’s instructions. One-step RT-qPCR was subsequently performed with the Luna Universal RT-qPCR Kit (New England Biolabs, #E3005L) according to the manufacturer’s instructions on a QuantStudio 5 Real-Time PCR System (Applied Biosystems) with SYBR Green detection chemistry. The thermal cycling conditions included initial denaturation and reverse transcription followed by amplification for 40 cycles. A melt curve analysis was performed to confirm amplification specificity. Custom primers were designed and synthesized by Integrated DNA Technologies (IDT). The following primers were used: human GPX4 forward (5’-GAGGCAAGACCGAAGTAAACTAC-3’) and reverse (5’- CCGAACTGGTTACACGGGAA-3’), human ACSL4 forward (5’- ACTGGCCGACCTAAGGGAG-3’) and reverse (5’-GCCAAAGGCAAGTAGCCAATA-3’), human FTH1 forward (5’-CGAGGTGGCCGAATCTTCC-3’) and reverse (5’- GTTTGTGCAGTTCCAGTAGTGA-3’), human FTL forward (5’- CAGCCTGGTCAATTTGTACCT-3’) and reverse (5’-GCCAATTCGCGGAAGAAGTG-3’), human SLC40A1 forward (5’-TGGATGGGTTCTCACTTCCTG-3’) and reverse (5’- GTCAATCCTTCGTATTGTGGCAT-3’), and human GAPDH forward (5’- ACAACTTTGGTATCGTGGAAGG-3’) and reverse (5’- GCCATCACGCCACAGTTTC-3’). Relative mRNA expression was determined using the 2^-ΔΔCT^ mathematical method and normalized to endogenous GAPDH.

### CRISPR-mediated generation of GPX4-knockout NCI-H295R cells

NCI-H295R GPX4 KO clones were stably generated using CRISPR genome-editing technology. Briefly, single guide RNA (sgRNA) targeting human *GPX4* (sequence: 5’- TAACCTGGACAAGTACCGGT-3’) was designed using the CRISPick sgRNA Designer (Broad Institute) and thereafter cloned into a lentiCRISPR v2-Blast vector plasmid (Addgene, #83480) in accordance with the manufacturer’s instructions. Lentiviral particles were produced and used to transduce approximately 5×10^5^ NCI-H295R cells. Transduced NCI-H295R cells were selected with 1 µg/mL blasticidin for 2 weeks, after which stable blasticidin-resistant cells were expanded and maintained under continued antibiotic selection to establish stable populations. GPX4 depletion was subsequently confirmed by Western blot analysis.

### *ABCB1*, *ABCC1*, and *ABCG2* overexpression assays

Stably-transfected human embryonic kidney (HEK-293) cells expressing *ABCB1*, *ABCC1*, or *ABCG2* were generated as previously described (56). Briefly, HEK293 cells were transfected with empty pcDNA3 vector (EV) (Invitrogen, CA, USA) or pcDNA3 vector containing full length *ABCB1*, *ABCC1*, or *ABCG2*. Transfectants were enforced with 2 mg/mL G418 and cultured in Eagle’s Minium Essential Medium (EMEM) (ATCC, 30-2003) supplemented with 10% FCS and 1% penicillin-streptomycin. For drug assays, transfected cells were seeded at a density of 5×10^3^ cells per well in opaque 96-well plates and allowed to attach overnight. Following overnight incubation, TR-107 was administered and cells were incubated for 72 hours. At the conclusion of the treatment period, CellTiter-Glo reagent was added to each treatment well and luminescence was quantified after a 100 ms read time using a Tecan Spark luminometer (Tecan US, Inc. Morrisville, NC).

### Western blotting

NCI-H295R, NCI-ACC48, and NCI-ACC60 PDOs were cultured as described above in tissue-culture 6-well plates until approximately 85% confluence. Cells and PDOs were then overlaid with fresh medium containing vehicle control DMSO or variable concentrations of TR- 107 for 48 hours. For basal protein expression, cells were lysed directly at 85% confluence. Cells and PDOs were dissociated into single cells and lysed with RIPA Lysis and Extraction Buffer (Thermo Scientific, #89900) supplemented with protease inhibitor (Thermo Scientific #78430) and 0.5 mM EDTA (Thermo Scientific, #R1021). Clarified protein lysate was obtained and quantified by Pierce BCA Protein Assay Kit (Thermo Scientific, #23227). Drug-treated or basal expression protein samples were prepared by mixing 25 µg of respective protein with 1x Bolt Sample Reducing Agent (10x) (Thermo Scientific, #B0009), RIPA buffer, and 1x Bolt LDS Sample Buffer (4x) (Invitrogen, #B0007). Prepared protein samples were denatured at 95°C for 5 minutes and separated using NuPAGE Bis-Tris Mini Protein Gels, 4-12%, 1.0-1.5 mm (Invitrogen, #NP0323BOX), transferred to nitrocellulose membranes (Invitrogen, #IB23001), and blocked for 1 hour with 5% weight/volume milk in 1x TBS-T solution. Following blocking, membranes were incubated with primary antibodies overnight at 4°C. The next day, membranes were washed with 1x TBS-T solution and then incubated with horseradish-peroxidase (HRP)- conjugated secondary antibodies in 5% milk in 1x TBS-T solution for 1 hour at room temperature. Signals were detected using the SuperSignal West Pico PLUS Chemiluminescent Substrate (Thermo Scientific, #34580) and imaged using the ChemiDoc MP Imaging System (Bio-Rad).

Primary and secondary antibodies were as follows: COX IV (3E11) (Cell Signaling Technology, #4850, rabbit, monoclonal, 1:1000 dilution), cytochrome c (136F3) (Cell Signaling Technology, #4280, rabbit, monoclonal, 1:1000 dilution), HSP60 (D6F1) (Cell Signaling Technology, #12165, rabbit, monoclonal, 1:1000 dilution), PHB1 (Cell Signaling Technology, #2426, rabbit, polyclonal, 1:1000 dilution), pyruvate dehydrogenase (C54G1) (Cell Signaling Technology, #3205, rabbit, monoclonal, 1:1000 dilution), SDHA (D6J9M) (Cell Signaling Technology, #11998, rabbit, monoclonal, 1:1000 dilution), SOD1 (71G8) (Cell Signaling Technology, #4266, mouse, monoclonal, 1:1000 dilution), SF-1 (A-1) (Santa Cruz Biotechnology, #sc-393592, mouse, monoclonal, 1:1000 dilution), OxPhos Rodent WB Antibody Cocktail (Invitrogen, #45-8099, mouse, polyclonal, 1:5000 dilution), CLPP (ABclonal, #A3214, rabbit, monoclonal, 1:1000 dilution), GPX4 (Cell Signaling Technology, #52455, rabbit, polyclonal, 1:1000 dilution), NCOA4 (E8H8Z) (Cell Signaling Technology, #66849, rabbit, monoclonal, 1:1000 dilution), KEAP1 (D6B12) (Cell Signaling Technology, #8047, rabbit, monoclonal, 1:1000 dilution), NRF2 (D1Z9C) (Cell Signaling Technology, #12721, rabbit, monoclonal, 1:1000 dilution), 4F2hc/SLC3A2 (D3F9D) (Cell Signaling Technology, #47213, rabbit, monoclonal, 1:1000 dilution), FTH1 (D1D4) (Cell Signaling Technology, #4393, rabbit, monoclonal, 1:1000 dilution), xCT/SLC7A11 (D2M7A) (Cell Signaling Technology, #12691, rabbit, monoclonal, 1:1000 dilution), DMT1/SLC11A2 (D3V8G) (Cell Signaling Technology, #15083, rabbit, monoclonal, 1:1000 dilution), AIFM2/FSP1 (Cell Signaling Technology, #24972, rabbit, polyclonal, 1:1000 dilution), ACSL4 (F6T3Z) (Cell Signaling Technology, #38493, rabbit, monoclonal, 1:1000 dilution), cleaved caspase-7 (Asp198) (D6H1) (Cell Signaling Technology, #8438, rabbit, monoclonal, 1:1000 dilution), cleaved caspase-9 (Asp330) (D2D4) (Cell Signaling Technology, #7237, rabbit, polyclonal, 1:1000 dilution), phospho-histone H2A.X (Ser139) (20E3) (Cell Signaling Technology, #9718, rabbit, monoclonal, 1:1000 dilution), ATF-4 (D4B8) (Cell Signaling Technology, #11815, rabbit, monoclonal, 1:1000 dilution), MDR1/ABCB1 (E1Y7B) (Cell Signaling Technology, #13342, rabbit, monoclonal, 1:1000 dilution), ABCG2 (D5V2K) (Cell Signaling Technology, #42078, rabbit, monoclonal, 1:1000 dilution), IGF-1R beta (D23H3) (Cell Signaling Technology, #9750, rabbit, monoclonal), IGF-II/IGF-2 (HL1982) (Novus Biologicals, #NBP3-25520, rabbit, monoclonal), histone H3 (1B1B2) (Cell Signaling Technology, #14269, mouse, monoclonal, 1:1000 dilution), HRP-linked anti-mouse IgG (Cell Signaling Technology, #7076), and HRP-linked anti-rabbit IgG (Cell Signaling Technology, #7074).

### Statistical analyses

All statistical tests were performed using GraphPad Prism 11.0 (RRID: SCR_002798). Comparisons among three or more groups were conducted using ordinary one-way analysis of variance (ANOVA) or two-way ANOVA with Tukey’s multiple comparisons test as appropriate for the experimental design. Statistically significant *p*-values are as follows: *P* < 0.05 (*), *P* < 0.01 (**), *P* < 0.001 (***), *P* < 0.0001 (****). For violin plots, horizonal dashed lines represent the median, while lower and upper dotted lines correspond to the first and third quartiles, respectively.

## Results

### TR-107 achieves potent, dose-dependent antitumor activity in ACC cell lines and short-term patient-derived organoid models

To evaluate the cytotoxic and inhibitory effects of TR-107 in ACC, we employed a heterogeneous panel of preclinical models comprising established ACC cell lines and novel short-term ACC patient-derived organoids (NCI-ACC PDOs) that we previously reported (49). ClpP expression was confirmed in representative ACC models (Supplementary Fig. S1A). ACC cells and PDOs were exposed to increasing concentrations of TR-107 for 72 hours, and thereafter viability was quantified and normalized to the vehicle control populations. TR-107 reduced viability across all ACC preclinical models in a dose-dependent manner (Fig. 1B). NCI-H295R cells were highly sensitive to TR-107, with a mean calculated half maximal inhibitory concentration (IC_50_) of 15.03 nmol/L (95% CI: 13.44 to 16.83 nmol/L), whereas mACC3 cells were comparatively less sensitive with a mean calculated IC_50_ of 99.65 nmol/L (95% CI: 89.29 to 116.00 nmol/L). Drug sensitivity varied broadly among NCI-ACC PDOs, with IC_50_ values spanning approximately 7.3-fold (11.32 nmol/L to 82.11 nmol/L), underscoring the intertumoral heterogeneity of patient-derived tumor samples. NCI-ACC PDOs exhibited mean calculated IC_50_ values of 28.00 nmol/L (95% CI: 15.45 to 53.02 nmol/L) for NCI-ACC45, 62.10 nmol/L (95% CI: 45.89 to 85.21 nmol/L) for NCI-ACC48, 13.80 nmol/L (95% CI: 9.507 to 20.08 nmol/L) for NCI-ACC56, 25.26 nmol/L (95% CI: 16.76 to 37.94 nmol/L) for NCI-ACC60, 11.32 nmol/L (95% CI: 7.100 to 17.81 nmol/L) for NCI-ACC74, 31.93 nmol/L (95% CI: 20.61 to 49.79 nmol/L) for NCI-ACC88, and 82.11 nmol/L (95% CI: for 56.10 to 122.9 nmol/L) for NCI- ACC95.

To assess the antiproliferative effects of TR-107, mACC3, NCI-H295R, and NCI-ACC60 PDOs were cultured with increasing concentrations of TR-107 (10 nmol/L to 80 nmol/L) or vehicle control and imaged by the Incucyte S3 Live-Cell Analysis Instrument. TR-107 treatment resulted in progressive and dose-dependent reduction in longitudinal cell confluence (Fig. 1C). Notably, treatment with 20 nmol/L of TR-107 was sufficient to markedly suppress proliferation across all evaluated models. Consistent with these findings, clonogenic assays demonstrated reductions in colony-forming capacity of NCI-H295R and mACC3 cells following TR-107 treatment, with 20 nmol/L inducing a sustained impairment of long-term proliferative capacity across both models (*P* < 0.0001). Moreover, colony formation was effectively abolished at 40 nmol/L and 80 nmol/L TR-107 (*P* < 0.0001) (Fig. 1D and E). Scratch wound assays further revealed a dose-dependent inhibition of tumor migratory potential following TR-107 treatment. Vehicle-treated NCI-H295R cells displayed progressive wound closure, with wound density approaching approximately 24-25% by 48 hours, indicative of intact migratory and proliferative capacity. Treatment with 1 nmol/L TR-107 did not measurably impair cell migration or proliferation, while 10 nmol/L TR-107 produced modest reductions in wound density (*P* < 0.05) but did not elicit a pronounced antiproliferative effect. In contrast, treatment with 100 nmol/L and 1 µmol/L resulted in significant reductions in wound density relative to vehicle control (*P* < 0.001 and *P* < 0.0001, respectively), and wound closure was significantly impaired at 1 µmol/L (*P* < 0.0001) (Fig. 1F and G). To better characterize cellular responses to TR-107, flow cytometric analyses were conducted on NCI-H295R cells following 48 hours of treatment to quantify proliferative and cytotoxic activity. EdU incorporation decreased in a dose-dependent manner, consistent with impaired DNA synthesis and concomitant G0/G1 or G2/M arrest (Fig. 1H; Supplementary Figure S1B). Annexin V/Propidium Iodide (PI) staining indicated dose-dependent increases in apoptotic cell fractions, with Annexin V^+^ cells increasing from 4.00% to 7.69% and Annexin V^+^/PI^+^ cells increasing from 6.31% to 18.60% relative to vehicle-treated cells (Fig. 1I; Supplementary Fig. S1C). Overall, these data suggest TR-107 exerts strong dose-dependent inhibitory effects on ACC cell proliferation, DNA synthesis, long-term clonogenic potential, and migratory capacity.

To further evaluate the antitumor efficacy of TR-107 in more clinically and physiologically relevant models, we generated NCI-ACC48 and NCI-ACC60 PDOs (Fig. 1J). Subsequent treatment over 48 hours with increasing concentrations of TR-107 revealed dose-dependent increases in Image-IT DEAD Green fluorescence across both PDO models, consistent with increased cell death and compromised membrane integrity compared to vehicle-treated controls (Fig. 1K). Quantitative analyses of the fluorescence at 488 nm demonstrated significant increases in the relative corrected total cell fluorescence (CTCF) ratio following increasing concentrations of TR-107 across both PDO models (*P* < 0.0001) (Fig. 1L). Generated NCI- ACC48 and NCI-ACC60 were immunostained for well-established ACC diagnostic and prognostic biomarkers, including steroidogenic factor 1 (SF-1), inhibin-α, melan-A, and β- catenin, and demonstrated positive staining (Fig. 1M) (57). In addition, hematoxylin and eosin (H&E) staining of NCI-ACC60 further corroborate tumor-like histopathological features (Supplementary Fig. S1D). Finally, to assess TR-107 selectivity, human embryonic kidney cell line HEK293T and normal human kidney organoid NCI-NKO were treated with increasing concentrations of TR-107 for 48 hours. Quantified viability data revealed significantly higher viability at equivalent doses relative to the ACC cell line NCI-H295R, thereby indicating selective cytotoxicity toward ACC (Fig. 1N). Collectively, these findings identify the highly potent and selective *in vitro* antitumor efficacy of TR-107 against variable ACC preclinical models.

### TR-107 disrupts mitochondrial function and attenuates steroid hormone secretion

Adrenocortical tumors arise from the highly steroidogenic adrenal cortex, a tissue with intrinsic dependence on mitochondrial metabolism to facilitate steroid hormone biosynthesis. Consequently, the pathophysiology of ACC is characterized by a distinctive metabolic architecture that drives aberrant steroidogenesis and extensive metabolic rewiring (2,6,58). To further characterize the metabolic landscape of ACC, we performed single-sample gene set enrichment analysis (ssGSEA) of curated metabolic pathway signatures across resected primary and metastatic ACC tissue specimens from the NCI-ACC institutional tumor tissue repository (*n* = 48 samples) and the publicly available The Cancer Genome Atlas (TCGA) ACC tumor cohort (*n* = 78 samples). OXPHOS exhibited marked enrichment across both cohorts irrespective of tumor MKI67 expression or variability in other metabolic and proliferative pathway signatures, thereby underscoring a conserved mitochondrial metabolic program in ACC (Fig. 2A).

**Figure 2.**
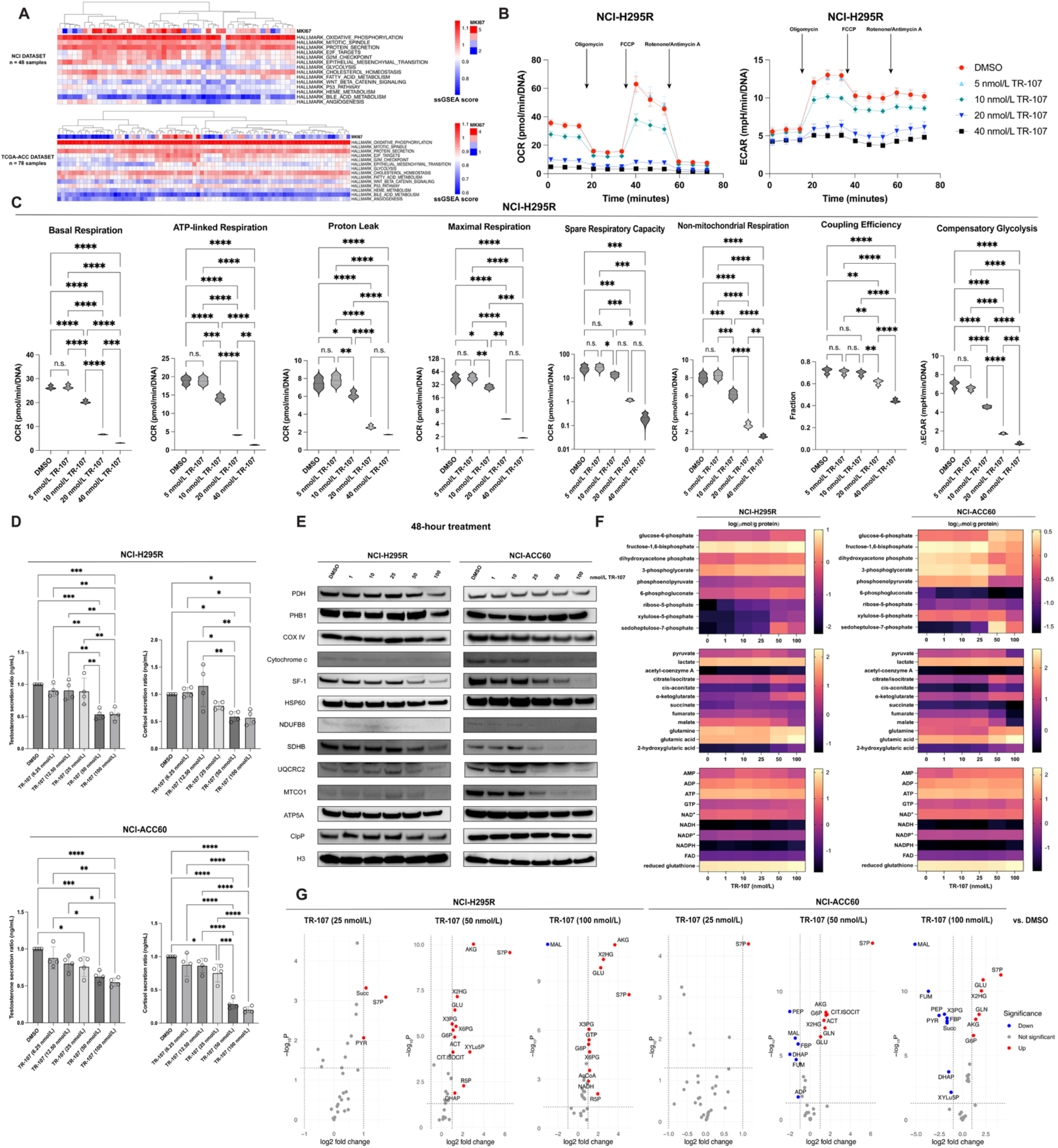
TR-107 suppresses mitochondrial respiratory function, attenuates sterodogenesis, and induces global bioenergetic collapse. **A,** Single-sample gene set enrichment analysis (ssGSEA) heatmap depicting enrichment of metabolic pathway signatures in NCI-ACC (*n* = 48 tumor specimens) and TCGA-ACC (*n* = 78 tumor specimens). MKI67, Ki-67 proliferation index. **B,** Oxygen consumption rate (OCR) (pmol/min) and extracellular acidification rate (ECAR) (mpH/min) measurements over time normalized to DNA in NCI-H295R cells following variable concentrations of TR-107. Data reported as mean ± standard deviation, *n* = 4 technical replicates. **C,** Quantification of mitochondrial function markers in NCI-H295R cells following TR-107 treatment. **D,** ELISA-based quantification of testosterone and cortisol secretion ratios in ng/mL in NCI-H295R cells and NCI-ACC60 PDOs following increasing concentrations of TR-107 treatment. Data reported as mean ± standard deviation, *n* = 4 biological replicates. **E,** Immunoblot analysis of essential mitochondrial proteins following 48-hour TR-107 treatment. PDH, pyruvate dehydrogenase; PHB1, prohibitin-1; COX IV, cytochrome c oxidase subunit IV; SF-1, steroidogenic factor-1; HSP60, heat shock protein 60; NDUFB8, NADH:ubiquinone oxidoreductase subunit B8; SDHB, succinate dehydrogenase complex iron-sulfur subunit B; UQCRC2, ubiquinol-cytochrome c reductase core protein 2; MTCO1, mitochondrially encoded cytochrome c oxidase I; ATP5A, ATP synthase F1 subunit alpha; ClpP, caseinolytic mitochondrial matrix peptidase proteolytic subunit; H3, histone H3. **F** and **G,** Targeted metabolomic profiling of NCI-H295R cells following TR-107 treatment using stable isotope dilution LC-MS/MS based multiple reaction monitoring (MRM) assays (**F**) and comparison across treatment groups (**G**), *n* = 3 biological replicates. n.s., not significant; *, *P* < 0.05; **, *P* < 0.01; ***, *P* < 0.001; ****, *P* < 0.0001.

Additional pathway analyses across both cohorts demonstrated that ACC tumors exhibiting elevated OXPHOS enrichment signatures were strongly associated with increased fatty acid metabolism (Pearson’s *r* = 0.52418, *P* = 0.00013, and Pearson’s *r* = 0.74834, *P* = 3.4×10^-15^, respectively) and moderately associated with cholesterol metabolism (Pearson’s *r* = 0.24799, *P* = 0.0089, and Pearson’s *r* = 0.37582, *P* = 7.0×10^-4^, respectively), suggesting a bioenergetic phenotype driven by coordinated mitochondrial respiration and lipid metabolic activity (Supplementary Fig. S1E). Consistent with the steroidogenic phenotype of ACC, OXPHOS-high adrenocortical tumors exhibited increased mRNA expression of essential steroidogenic modulators including STAR, CYP11A1, CYP21A2, CYP11B1, and NR5A1, among both NCI-ACC and TCGA-ACC cohorts (Supplementary Fig. S1F and G). This associated transcriptional enrichment suggests a relationship between mitochondrial respiratory state and steroidogenic activity, thereby highlighting an integrated metabolic-steroidogenic program that may contribute to the defining features of ACC pathophysiology.

Previous investigation of TR-107 has demonstrated robust perturbation of mitochondrial respiration in breast cancer and colorectal cancer cells (11,12). To assess whether TR-107 similarly impairs mitochondrial function in ACC, NCI-H295R cells were treated with TR-107 for 48 hours, and mitochondrial respiration was subsequently analyzed using the Seahorse XFe96/XF96 Extracellular Flux Analyzer (Agilent Technologies). We observed pronounced, dose-dependent reductions in oxygen consumption rate (OCR) following TR-107 treatment (Fig. 2B). Treatment with 5 nmol/L TR-107 did not significantly alter basal respiration or other parameters of mitochondrial function relative to the vehicle control. In contrast, 10 nmol/L TR- 107 induced modest but significant reductions in basal respiration, accompanied by significant decreases in ATP-linked respiration, proton leak, maximal respiration, and non-mitochondrial respiration. Moreover, 10 nmol/L TR-107 moderately reduced the extracellular acidification rate (ECAR) and crude measures of compensatory glycolysis. Conversely, TR-107 induced near-complete abolishment of basal respiration, ATP-linked respiration, proton leak, maximal respiration, spare respiratory capacity, non-mitochondrial respiration, and significant reductions in coupling efficiency at ≥ 20 nmol/L. Furthermore, NCI-H295R cells treated with 20 and 40 nmol/L of TR-107 failed to increase ECAR, indicating an inability to compensate for mitochondrial dysfunction through glycolysis (Fig. 2C). In addition, ELISA-based quantification of testosterone and cortisol secretion in NCI-H295R cells and NCI-ACC60 PDOs revealed that increasing concentrations of TR-107 substantially attenuates steroid hormone production and secretion (Fig. 2D). Analyses of protein expression in TR-107-treated NCI-H295R and NCI- ACC60 PDO cells revealed dose-dependent depletion of mitochondrial respiratory chain components, including subunits of complex I-IV, mitochondrial regulatory proteins such as cytochrome c and HSP60, and steroidogenesis-associated transcription factor SF-1 (Fig. 2E). Interestingly, pyruvate dehydrogenase expression showed depletion at high dose (100 nmol/L) TR-107, suggesting relative preservation of pyruvate oxidation and acetyl-CoA synthesis. In addition, ATP5A expression remained relatively stable, suggesting some preservation of the ATP synthase complex despite TR-107-induced metabolic remodeling, while increased expression of scaffold protein prohibitin 1 suggests compensatory stabilization of mitochondrial cristae architecture. Ultimately, these findings suggest that TR-107 induces robust disruption of essential respiratory function and steroidogenic capacity in ACC.

To further define the metabolic consequences of TR-107-mediated mitochondrial disruption, we performed targeted metabolomic profiling of TR-107-treated NCI-H295R and NCI-ACC60 PDOs. High-dose TR-107 (50 nmol/L and 100 nmol/L) reduced cellular ATP levels while increasing NADH abundance, consistent with impaired electron transfer and accumulation of reduced cofactors (59). In addition, significant reductions in fumarate and malate levels support disruption of TCA cycle metabolism and impaired electron transport chain function, including perturbation of complex II. While marked depletion of critical canonical glycolytic intermediates, including phosphoenolpyruvate, pyruvate, and fructose-1,6-bisphosphate suggest impaired glycolytic flux, the dramatic accumulation of ribose-5-phosphate, xylulose-5- phosphate, and sedoheptulose-7-phosphate across both models suggests attempted compensation through the pentose phosphate pathway to refuel glycolysis (60). Moreover, accumulation of α- ketoglutarate, glutamic acid, and glutathione together in the setting of depleted downstream TCA cycle intermediates suggests altered glutamine metabolism in response to respiratory impairment, potentially reflecting enhanced glutaminolysis or amino acid catabolism (Fig. 2F and G) (61). Taken together, these metabolic perturbations reveal that TR-107 induces a profound bioenergetic crisis in ACC, characterized by impaired cellular respiration, altered metabolic profile, and failure to activate compensatory metabolic adaptations.

### TR-107 drives ferroptotic cell death through ROS accumulation, lipid peroxidation, and suppressed antioxidant defense

Ferroptosis is an iron-dependent form of regulated death driven by reactive oxygen species (ROS) and distinctly characterized by marked intracellular iron accumulation and excessive lipid peroxidation (62–66). The multifaceted regulatory role of mitochondria in bioenergetics, redox homeostasis, iron metabolism, and cell fate positions these organelles as central modulators of ferroptotic vulnerability, with mitochondrial dysfunction promoting oxidative stress and governing pro-ferroptotic signaling cascades (64,65,67). ssGSEA-based assessment of the NCI-ACC institutional tumor tissue repository and the TCGA-ACC tumor cohorts revealed that OXPHOS-high adrenocortical tumors exhibited increased enrichment of ROS-associated genomic signatures (Pearson’s *r* = 0.39462, *P* = 0.0055, and Pearson’s *r* = 0.27266, *P* = 0.016, respectively), suggesting an intrinsically heightened oxidative state that may confer predisposed susceptibility to ferroptotic signaling pathways (Supplementary Fig. S2A). Previous studies have demonstrated that TR-107 induces ferroptotic cell death in colorectal cancer cells (11); however, the mechanisms mediating this response are not fully elucidated.

To better elucidate this ferroptotic response, we investigated whether TR-107 elicited similar effects in ACC models. We observed that co-treatment of NCI-H295R and NCI-ACC60 PDO cells with 10 µmol/L of small-molecule ferroptosis inhibitors liproxstatin-1 and ferrostatin- 1 significantly attenuated 25 nmol/L and 50 nmol/L TR-107 cytotoxicity (*P* < 0.0001) across both models (Fig. 3A). Corroborating these results, TR-107 induced dose-dependent increases in intracellular 2′,7′-dichlorofluorescein (DCF) fluorescence intensity in both NCI-H295R and NCI- ACC60 PDO cells, indicative of elevated ROS accumulation and enhanced oxidative stress. Notably, DCF fluorescence intensity declined with high doses of TR-107 (50 nmol/L and 100 nmol/L), likely reflecting extensive cytotoxicity and reduced viable cells available for ROS detection (Fig. 3B). Furthermore, TR-107-treated NCI-H295R and NCI-ACC60 cells exhibited significant dose-dependent increases in malondialdehyde, a principal aldehyde byproduct formed from ROS-mediated oxidation of membrane polyunsaturated fatty acids (Fig. 3C). Quantitative RT-PCR profiling of ferroptosis-associated genes in NCI-H295R and NCI-ACC60 PDO cells following 24-hour TR-107 treatment demonstrated a shift toward ferroptosis-permissive transcriptional state, with reduced expression of glutathione peroxidase 4 (GPX4) and ferritin components ferritin light chain (FTL) and ferritin heavy chain (FTH1), alongside increased expression of acyl-CoA synthetase long chain family member 4 (ACSL4) and ferroportin (SLC40A1) (Fig. 3D). These data indicate that TR-107 induces concerted transcriptional remodeling of ferroptosis-regulatory pathways that impairs antioxidant defense, alters cellular iron-handling, and enables a ferroptosis-permissive state.

**Figure 3.**
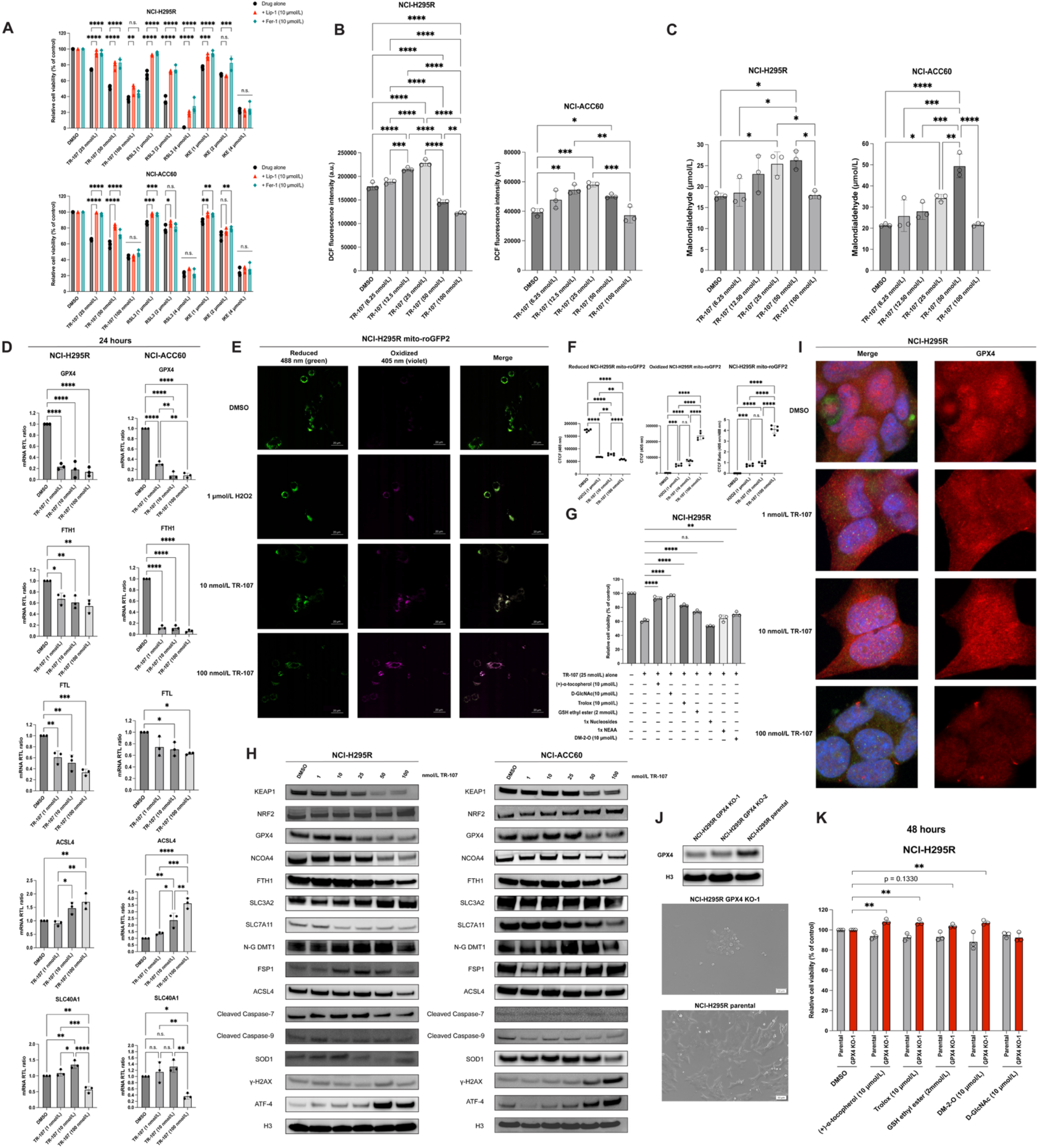
TR-107 shifts the ferroptotic rheostat toward a ferroptosis-permissive state by promoting oxidative stress, lipid peroxidation, and impaired cellular antioxidant capacity. **A,** Cell viability of NCI-H295R cells and NCI-ACC60 PDOs following 48-hour treatment with variable concentrations of TR-107 in the presence or absence of ferroptosis inhibitors liproxstatin-1 (Lip- 1; 10 µmol/L) and ferrostatin-1 (Fer-1; 10 µmol/L). Cell viability was normalized to vehicle- treated controls and expressed as a percentage of vehicle controls. RSL3, (1S,3R)-RSL3; IKE, imidazole ketone erastin. **B** and **C,** Measurement of intracellular reactive oxygen species (ROS) by 2′,7′-dichlorofluorescein (DCF) fluorescence intensity (**B**) and cellular malondialdehyde concentrations in µmol/L (**C**) in NCI-H295R cells and NCI-ACC60 PDOs treated with increasing concentrations of TR-107 for 48 hours. **D,** RT-qPCR analysis of ferroptosis-associated genes in NCI-H295R cells and NCI-ACC60 PDOs following 24-hour TR-107 treatment. *Y*-axis: mRNA relative transcription level ratio. Data reported as mean ± standard deviation, *n* = 3 technical replicates. GPX4, glutathione peroxidase 4; FTH1, ferritin heavy chain 1; FTL, ferritin light chain; ACSL4, acyl-CoA synthetase long-chain family member 4; SLC40A1, solute carrier family 40 member 1. **E** and **F,** Visualization (**E**) and fluorescence intensity quantification (**F**) of mitochondrial oxidative stress using NCI-H295R mito-roGFP2 cells following TR-107 treatment. Green fluorescence at 488 nm indicates a global reduced mitochondrial state; violet fluorescence at 405 nm indicates a global oxidized mitochondrial state. Quantification was performed from five independent fluorescence measurements per condition. **G,** Co-treatment of NCI-H295R cells with TR-107 and the indicated pharmacologic antioxidants for 48 hours. Cell viability was normalized to vehicle-treated controls and expressed as a percentage of vehicle controls. D-GlcNAc, N-Acetyl-D-glucosamine; Trolox, (6-hydroxy-2,5,7,8-tetramethylchroman- 2-carboxylic acid); NEAA, non-essential amino acids; DM-2-O, dimethyl-2-oxoglutarate. **H,** Immunoblot analysis of variable ferroptosis-associated proteins following 48-hour TR-107 treatment in NCI-H295R cells and NCI-ACC60 PDOs. KEAP1, Kelch-like ECH-associated protein 1; NRF2, nuclear factor erythroid 2-related factor; GPX4, glutathione peroxidase 4; NCOA4, nuclear receptor coactivator 4; FTH1, ferritin heavy chain 1; SLC3A2, solute carrier family 3 member 2; SLC7A11, solute carrier family 7 member 11; N-G DMT1, non-glycosylated divalent metal transporter 1; FSP1, ferroptosis suppressor protein 1; ACSL4, acyl-CoA synthetase long-chain family member 4; SOD1, superoxide dismutase 1; γ-H2AX, gamma-H2A histone family member X; ATF-4, activating transcription factor 4; H3, histone H3 **I,** Immunofluorescent staining of GPX4 protein in NCI-H295R cells following 48-hour TR-107 treatment. Red, GPX4; green, phalloidin; blue, DAPI. **J,** Immunoblot validation of NCI-H295R GPX4 KO cells and representative morphological images. **K,** 48-hour treatment of NCI-H295R GPX4 KO cells with pharmacologic antioxidants. Cell viability was normalized to vehicle- treated controls and expressed as a percentage of vehicle controls. Data reported as mean ± standard deviation, *n* = 3 technical replicates unless otherwise specified. n.s., not significant; *, *P* < 0.05; **, *P* < 0.01; ***, *P* < 0.001; ****, *P* < 0.0001.

To further delineate the contribution of oxidative stress to TR-107 cytotoxicity, we assessed changes in intracellular redox homeostasis dynamics using a redox-sensitive mito- roGFP2 reporter system in NCI-H295R cells. Exposure to increasing concentrations of TR-107 for 48 hours resulted in a significant, dose-dependent increases (*P* < 0.0001) in the oxidized/reduced CTCF ratio in NCI-H295R mito-roGFP2 cells compared to vehicle control, reflecting a pronounced shift toward a more oxidized mitochondrial redox environment. Notably, high dose TR-107 (100 nmol/L) produced a markedly greater oxidative shift that exceeded the response observed with oxidative stress positive control 1 µmol/L H_2_O_2_ (Fig. 3E and F) We next evaluated whether restoration of antioxidant capacity could rescue TR-107-mediated cytotoxicity. NCI-H295R cells co-treated with 25 nmol/L of TR-107 and a panel of pharmacologic antioxidants demonstrated significant restoration (*P* < 0.01 and *P* < 0.0001) of relative cell viability compared to TR-107 monotherapy alone. Strikingly, antioxidants which directly scavenge lipid peroxyl radicals within membranes, including (+)-α-tocopherol and related α-tocopherol analog Trolox, as well as indirect ROS suppressors such as N-acetyl-D- glucosamine, produced the strongest protective effects (Fig. 3G). Given the evidence of TR-107- induced lipid peroxidation and ROS accumulation, we next sought to define the molecular mechanisms underlying the ferroptotic response in ACC. Immunoblotting of NCI-H295R cells and NCI-ACC60 PDOs following 48-hour TR-107 treatment subsequently revealed dynamic dose-dependent modulation of proteins governing the ferroptotic rheostat (Fig. 3H) (11,62,63). In both models, TR-107 elicited concerted depletion of critical ferroptosis defense mediators, including GPX4, FTH1 SLC7A11, and SOD1, thereby compromising peroxide detoxification and scavenging capacity, iron homeostasis, and redox control. Conversely, NRF2 and FSP1 expression were elevated, suggesting activation of an adaptive NRF2-mediated antioxidant stress response to mitigate mounting oxidative stress. In parallel, TR-107 induced modest ACSL4 protein expression, while increased γ-H2AX at higher TR-107 concentrations demonstrated accumulation of DNA damage associated with severe oxidative stress. Finally, ATF-4 upregulation further reflects activation of the canonical integrated stress response, a well- established consequence of mitochondrial perturbation and dysregulated mitochondrial homeostasis (33).

Immunofluorescent analysis of 48-hour TR-107-treated NCI-H295R cells further validated a dose-dependent reduction in GPX4 protein abundance (Fig. 3I). Given the indispensable role of reduced glutathione (GSH) as the reducing co-substrate required for GPX4- mediated lipid hydroperoxide detoxification and regeneration of active GPX4, the dramatic accumulation of GSH observed by metabolomic profiling in both ACC models is consistent with impaired GPX4 activity (Fig. 2G). To establish the functional relevance of GPX4 loss, we next generated NCI-H295R GPX4 KO cells exhibited striking morphological alterations and diminished cellular viability, consistent with profound oxidative injury (Fig. 3J; Supplementary Fig. S2B). Importantly, supplementation with a similar panel of pharmacologic antioxidants stimulates basal NCI-H295R GPX4 KO cell viability relative to vehicle control, thus implying that selective GPX4 depletion recapitulates the redox vulnerabilities observed following TR-107 treatment (Fig. 3K). Collectively, these results validate TR-107 as a potent ferroptosis-inducing compound that precipitates mitochondrial redox collapse, elicits robust oxidative stress, attenuates cellular antioxidant mechanisms, and establishes a ferroptosis-permissive state in ACC models.

### TR-107 is not a substrate of the ABCB1 drug efflux transporter

ATP-binding cassette (ABC) transporter proteins are ubiquitously expressed in mammalian cell membranes and primarily function to secrete xenobiotic and cytotoxic compounds. ABC transporters have been identified as key mediators of multidrug resistance in tumors, with ABCB1 (MDR1/P-glycoprotein) and ABCG2 (MXR/BCRP) efflux transporters integral to resistance against a broad spectrum of clinically active anticancer drugs, including doxorubicin, etoposide, imatinib, paclitaxel, cisplatin, topotecan, and carboplatin (68,69). The ABCB1 efflux transporter has particularly been implicated in clinical multidrug resistance and increased adverse toxicities in the context of ACC (69). Western blot analysis confirmed the differential expression of ABCB1 and ABCG2 in NCI-H295R and representative PDOs NCI- ACC48 and NCI-ACC60, which themselves were generated from highly chemoresistant patient tumors (Table 1; Fig. 4A). Consistent with their clinical phenotypic profiles, NCI-ACC48 and NCI-ACC60 PDOs exhibited marked resistance to widely used chemotherapeutic agents that are established ABCB1 and ABCG2 substrates, as well as to the standard-of-care agent mitotane (Fig. 4B). In NCI-ACC48, the mean calculated IC_50_ values were 13.07 µmol/L (95% CI: 12.70 to 13.45 µmol/L) for mitotane and 24.19 µmol/L (95% CI: 20.08 to 29.41 µmol/L) for doxorubicin, while carboplatin, etoposide, and cisplatin did not reach measurable IC_50_ values at the concentrations tested. In NCI-ACC60, the mean calculated IC_50_ values were 15.88 µmol/L (95% CI: 15.36 to 16.42 µmol/L) for mitotane, 5.149 µmol/L (95% CI: 4.923 to 5.361 µmol/L) for doxorubicin, 2872 µmol/L (95% CI: 1001 to 19877 µmol/L) for carboplatin, 496.8 µmol/L (95% CI: 305.3 to 1048 µmol/L) for etoposide, and 339.3 µmol/L (95% CI: 194 to 1021 µmol/L) for cisplatin.

**Figure 4.**
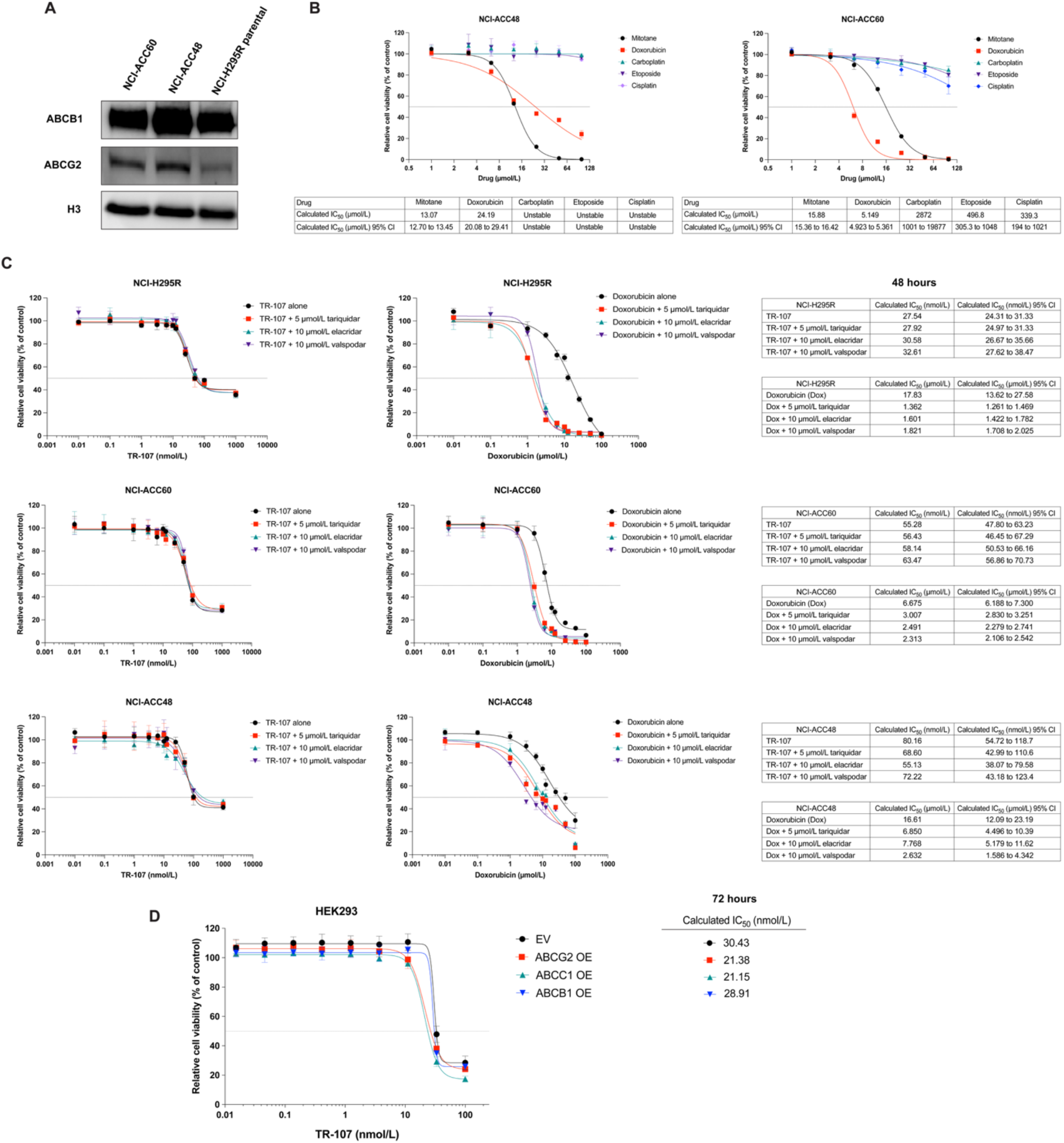
TR-107 is not a substrate of the ABCB1/MDR1/P-glycoprotein. **A,** Immunoblot validation of ABCB1 and ABCG2 protein expression in representative ACC models NCI-H295R, NCI- ACC48, and NCI-ACC60. H3, histone H3. **B,** Dose-response curves of variable chemotherapeutic agents following 48-hour treatment across NCI-ACC48 and NCI-ACC60 PDOs with corresponding IC_50_ values in µmol/L. Cell viability was normalized to vehicle-treated controls and expressed as a percentage of vehicle controls. **C,** Dose-response curves of TR-107 or doxorubicin co-treated with ABC transporter inhibitors tariquador (5 µmol/L), elacridar (10 µmol/L), or valspodar (10 µmol/L) for 48 hours across NCI-H295R, NCI-ACC48 PDOs, and NCI-ACC60 PDOs. Cell viability was normalized to vehicle-treated controls and expressed as a percentage of vehicle controls. Corresponding IC_50_ values were calculated and provided in nmol/L or µmol/L, as indicated. **D,** Dose-response curves of TR-107 following 72-hour treatment in HEK293 cells expressing empty vector, *ABCG2*, *ABCC1*, or *ABCB1*, with corresponding IC_50_ values provided. Cell viability was normalized to vehicle-treated controls and expressed as a percentage of vehicle controls. EV, empty vector. OE, overexpression. Data reported as mean ± standard deviation, *n* = 3 technical replicates.

To determine whether TR-107 is a substrate of the ABCB1or ABCG2 efflux transporters, TR-107 and doxorubicin were co-administered to NCI-H295R, NCI-ACC48, and NCI-ACC60 PDOs in the presence of highly potent and selective ABCB1 inhibitors elacridar (10 µmol/L) or valspodar (10 µmol/L), or the dual ABCB1/ABCG2 inhibitor tariquidar (5 µmol/L). Notably, inhibition of ABCB1 or ABCG2 alone did not significantly impact cell viability (Supplementary Fig. S2C). Across all evaluated models, co-treatment with variable ABCB1 or ABCG2 inhibitors did not produce a meaningful shift in the mean calculated IC_50_ for TR-107, with all calculated values remaining within overlapping 95% confidence intervals with respect to each model (Fig. 4C). In contrast, the established ABCB1/ABCG2 substrate doxorubicin displayed the expected increase in sensitivity upon co-treatment with ABCB1/ABCG2 inhibitors, thereby validating the functional activity of the inhibitors in these model systems. In addition, overexpression of ABCB1, ABCC1, or ABCG2 efflux transporters in HEK293T cells did not produce meaningful leftward shifts in the dose-response of TR-107 compared with empty vector controls (Fig. 4D). Overall, these findings provide compelling evidence that TR-107 is not a substrate of the ABCB1 or ABCG2 drug efflux transporters and is therefore not subject to ABCB1/ABCG2-mediated drug efflux.

### TR-107 synergizes with IGF-1R blockade to induce greater antitumor activity

Aberrations in the insulin-like growth factor 2/insulin-like growth factor 1 receptor (IGF- 2/IGF-1R) signaling axis are classical hallmarks of adrenocortical tumors, in which dysregulated *IGF2* expression drives pleiotropic proliferative effects through canonical Ras/Raf/MAPK/ERK and PI3K/Akt/mTOR signaling cascades, thereby orchestrating robust oncogenic processes including cell-cycle progression, metabolic reprogramming, evasion of apoptosis, and metastasis (70,71). Western blot analysis of NCI-H295R, NCI-ACC48, and NCI-ACC60 cells confirmed abundant IGF-1R and IGF-2 protein expression, while ELISA-based quantification further validated IGF-2 secretion, thereby supporting an active autocrine IGF-2/IGF-1R signaling axis across our evaluated ACC models (Fig. 5A and B). Interestingly, analyses of the NCI-ACC tumor tissue repository revealed that IGF2 expression was moderately correlated with high-OXPHOS signatures (Pearson’s *r* = 0.30029, *p* = 0.038) (Supplementary Fig. S2D).

**Figure 5.**
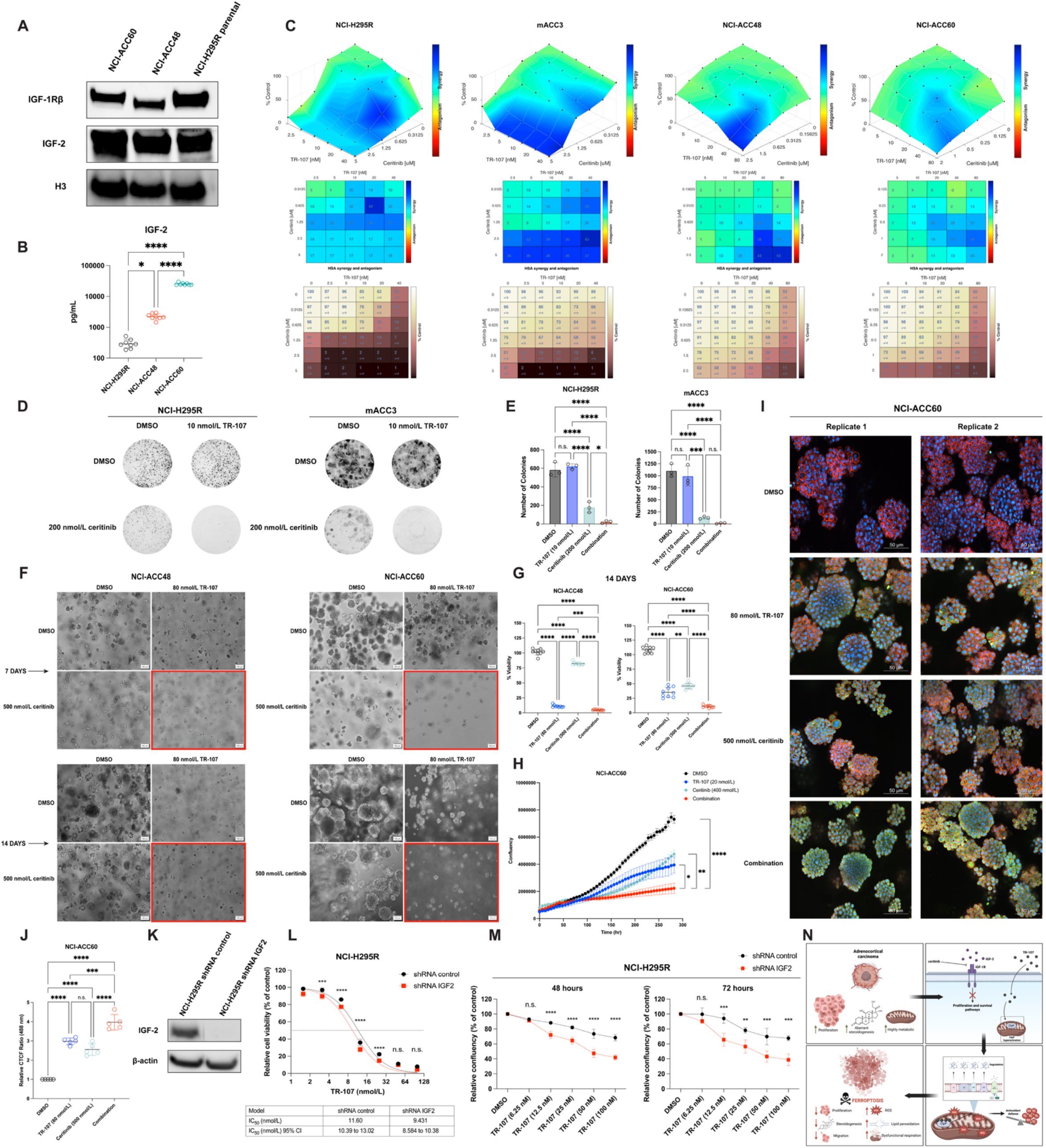
TR-107 synergizes with IGF-1R blockade in preclinical models of adrenocortical carcinoma. **A** and **B,** Immunoblot analysis of IGF-1Rβ and IGF-2 protein expression (**A**) and ELISA-based quantification of IGF-2 secretion (**B**) in representative ACC models NCI-H295R, NCI-ACC48, and NCI-ACC60, *n* = 4 biological replicates analyzed in duplicate. H3, histone H3. **C,** Synergistic dose-response matrices for TR-107 and ceritinib across NCI-H295R, mACC3, NCI- ACC48, and NCI-ACC60 models. Cell viability was normalized to vehicle-treated controls and expressed as a percentage of vehicle controls. **D** and **E,** Representative crystal violet staining (**D**) and quantification of NCI-H295R and mACC3 colony formation (**E**) following TR-107 plus ceritinib combinatory treatment, *n* = 3 technical replicates. **F** and **G,** Representative images (**F**) and quantification of viability (**G**) following TR-107 plus ceritinib combinatory treatment in NCI-ACC48 and NCI-ACC60 PDOs, *n* = 9 biological replicates. Images outlined in red indicate combinatory treatment. **H,** Longitudinal cell proliferation of NCI-ACC60 PDOs following TR- 107 plus ceritinib combinatory treatment. Data reported as mean ± SEM, *n* = 3 technical replicates. **I** and **J,** Representative fluorescent images of NCI-ACC60 PDOs following TR-107 plus ceritinib combinatory treatment (**I**) and quantification of green fluorescence (**J**). Green: Image IT-DEAD reagent; red: phalloidin; blue: DAPI. Quantification was performed from five independent fluorescence measurements per condition. For each measurement, fluorescence intensity values from two biological replicates were averaged and reported as individual data points. **K** and **L,** Immunoblot validation of NCI-H295R *IGF2* and control shRNA clones (**K**) and dose-response effect of TR-107 in NCI-H295R *IGF2* shRNA and NCI-H295R control shRNA cells (**L**). **M,** Analysis of proliferation in NCI-H295R *IGF2* shRNA and NCI-H295R control shRNA cells following TR-107 treatment. Data reported as mean ± SEM, *n* = 3 technical replicates. **N,** Schematic overview of the proposed mechanism of action of TR-107 and its augmentation with IGF-1R blockade in ACC. Data reported as mean ± standard deviation, *n* = 3 technical replicates unless otherwise specified. n.s., not significant; *, *P* < 0.05; **, *P* < 0.01; ***, *P* < 0.001; ****, *P* < 0.0001.

Given the essential role of the IGF-2/IGF-1R signaling axis in ACC, we next examined the combinatorial therapeutic potential of TR-107 with IGF-1R inhibitors. Ceritinib, an FDA-approved dual anaplastic lymphoma kinase (ALK) and IGF-1R inhibitor, demonstrated strong cytotoxic activity in the micromolar range and achieved the highest 48-hour synergistic activity in combination with TR-107 across the evaluated ACC models (Fig. 5C; Supplementary Fig. 2E). Notably, NCI-H295R *IGF1R* shRNA cells exhibited approximately 4-fold reduced sensitivity to 72-hour ceritinib treatment compared with control shRNA cells, with IC_50_ values of 3.990 µmol/L (95% CI: 3.652 to 4.549 µmol/L) versus 1.181 µmol/L (95% CI: 1.012 to 1.381 µmol/L), respectively (Supplementary Fig. S2F). Western blot analysis confirmed IGF-1R depletion in NCI-H295R *IGF1R* shRNA cells, thereby suggesting that ceritinib cytotoxicity is primarily mediated through the IGF-1R axis rather than through ALK inhibition in NCI-H295R cells (Supplementary Fig. S2G). Functional validation using clonogenic assays of treated NCI- H295R and mACC3 cells revealed that combined TR-107 (10 nmol/L) and ceritinib (200 nmol/L) treatment produced more pronounced reductions of long-term colony-forming capacity relative to either monotherapy or the vehicle control alone (Fig. 5D and E). Cytotoxicity assays further demonstrated enhanced antitumor activity following 72 hours of TR-107 and ceritinib co-treatment compared to either monotherapy or vehicle control alone across all evaluated ACC models (Supplementary Fig. 2H).

Representative images of NCI-ACC48 PDOs obtained at 7 and 14 days post-treatment demonstrated that 80 nmol/L TR-107 plus 500 nmol/L ceritinib treatment induced modest additional reductions in organoid growth relative to 80 nmol/L TR-107 alone, although both treatment conditions produced substantially greater growth suppression than either 500 nmol/L ceritinib or vehicle control alone. In contrast, representative images of NCI-ACC60 PDOs obtained at 7 and 14 days post-treatment revealed that 80 nmol/L TR-107 plus 500 nmol/L ceritinib treatment induced robust reductions in organoid growth compared with either monotherapy or the vehicle control alone (Fig. 5F). Consistent with these observations, quantified viability at 14 days post-treatment revealed significantly increased cytotoxic effect in combinatory treatment across NCI-ACC48 and NCI-ACC60 PDOs (Fig. 5G). However, in NCI- ACC48 PDOs, combination therapy had a modest effect compared to single agents.

Monotherapy with 80 nmol/L TR-107 or 500 nmol/L ceritinib reduced mean viability to 11.06% and 82.31%, respectively, with combination treatment producing a further reduction to 4.78%. In NCI-ACC60 PDOs, 80 nmol/L TR-107 or 500 nmol/L ceritinib monotherapy exhibited more modest effects to mean viability (35.17% and 46.18%, respectively), while combination treatment produced a substantial reduction of mean viability to 10.87%. Together, these data demonstrate augmented cytotoxicity in TR-107 plus ceritinib combinatory treatment across PDO models.

Consistent with the enhanced cytotoxic and inhibitory effect of TR-107 and ceritinib co-treatment in NCI-ACC60 PDOs, longitudinal live-cell proliferation analyses using the Incucyte S3 platform revealed sustained and significant suppression of organoid growth following 20 nmol/L TR-107 plus 400 nmol/L ceritinib combination therapy compared to either monotherapy (Fig. 5H). In addition, co-treatment with 80 nmol/L TR-107 and 500 nmol/L ceritinib profoundly induced Image-IT DEAD Green fluorescence, with quantitative analysis demonstrating a significant elevation (*P* < 0.001 and *P* < 0.0001) in the relative CTCF ratio compared to either monotherapy alone (Fig. 5I and J). To further evaluate the combinatorial potential of TR-107 with IGF-1R inhibition, stable NCI-H295R *IGF2* shRNA and NCI-H295R control shRNA cell lines were generated and validated by immunoblotting for IGF-2 (Fig. 5K). NCI-H295R *IGF2* shRNA cells demonstrated significantly higher sensitivity to 72-hour TR-107 treatment at low doses compared with NCI-H295R control shRNA cells (Fig. 5L). In addition, longitudinal monitoring of cell proliferation following treatment with increasing concentrations of TR-107 revealed significantly reduced confluency at equivalent doses (*P* < 0.01, *P* < 0.001, and *P* <0.0001) in NCI-H295R *IGF2* shRNA cells relative to NCI-H295R control shRNA cells, thereby indicating that IGF-2 depletion enhances the antiproliferative effects of TR-107 (Fig. 5M; Supplementary Fig. S2I). Together, these findings identify complementary metabolic and growth signaling vulnerabilities in ACC and demonstrate that concurrent targeting of mitochondrial respiratory function and the IGF-2/IGF-1R signaling axis potentiates antitumor activity beyond either strategy alone (Fig. 5N).

## Discussion

Adrenocortical carcinoma (ACC) is a rare and aggressive endocrine malignancy characterized by high recurrence rates, intrinsic chemoresistance, and limited effective systemic therapeutic options resulting in 5-year overall survival rates below 15% for patients with metastatic disease (1–4). Despite advances in our understanding of the molecular landscape of ACC, clinical responses to current standard of care agent, mitotane, as well as conventional chemotherapies are often transient and impaired by intrinsic or acquired drug resistance (2–4). The standard first-line systemic regimen for metastatic ACC, etoposide, doxorubicin, and cisplatin combined with mitotane (EDP-M), only achieves an objective response rate of approximately 23% with a median time to progression of approximately 5 months (1). These challenges reinforce the need to identify therapeutic strategies that better target biological dependencies unique to ACC. Herein, we identify TR-107 as a potent antitumor agent with dose-dependent, nanomolar activity across multiple preclinical models of ACC, consistent with prior investigation of TR-107 in triple-negative breast cancer, colorectal cancer, and glioblastoma cell lines (11,12,47). Our finding demonstrates that pharmacologic activation of ClpP by TR-107 produces robust antitumor activity across heterogenous ACC models, indicating that mitochondrial proteostasis represents a therapeutically actionable vulnerability in ACC. The observed antitumor activity across complementary preclinical systems with diverse genetic and phenotypic profiles suggests that the therapeutic potential of TR-107 is not confined to single molecular subtypes, thereby supporting its broader translational relevance in ACC. In addition to inducing cytotoxicity, TR-107 markedly impaired multiple hallmarks of tumor progression, including proliferation, clonogenic survival, DNA synthesis, and migratory capacity, suggesting that its antitumor effects extend beyond acute tumor cell killing. Most notably, TR-107 demonstrated preferential activity and high selectivity against ACC models while sparing normal renal epithelial cells and kidney-derived PDOs, implying a favorable therapeutic window and accentuating the potential for selective targeting of adrenocortical tumors.

Emerging evidence indicates an intrinsic reliance on mitochondria and respiratory machinery in ACC pathophysiology, suggesting that these tumors may be particularly vulnerable to respiratory disruption and subsequent bioenergetic collapse (2,6,10,16,58). Steroidogenesis is fundamentally a mitochondrial process, as the initial and rate-limiting step of steroid hormone biosynthesis, cholesterol side-chain cleavage to pregnenolone by CYP11A1, occurs in the inner mitochondrial membrane, and subsequent hydroxylation steps (e.g., 11β-hydroxylation by CYP11B1) also require functional mitochondrial respiratory chain activity (72) Notably, while our transcriptomic analyses demonstrate conserved OXPHOS enrichment in ACC tumors, prior proteomic studies have identified decreased expression of mitochondrial respiratory complex I and IV subunits in ACC compared with benign adrenocortical adenomas, potentially reflecting increased turnover or degradation of respiratory chain components in the setting of heightened metabolic demand (73). This apparent discrepancy between transcriptomic upregulation and proteomic depletion of OXPHOS components may indicate compensatory transcriptional responses to protein-level deficiencies, further underscoring the metabolic vulnerability of ACC to agents that disrupt mitochondrial proteostasis. ClpP-mediated proteolysis of OXPHOS respiratory components therefore represents a promising therapeutic strategy to expand precision medicine approaches that exploit such metabolic dependencies (10,11,26). Accordingly, metabolomic analyses following TR-107 treatment revealed broad and highly nuanced metabolic remodeling and imbalanced bioenergetic homeostasis across both NCI-H295R cell line and NCI- ACC60 PDO models. The observed metabolic alterations ultimately converge on impaired basal and ATP-linked respiration through depletion of critical electron transfer mediators, namely complexes I-IV and cytochrome c. In addition, the depletion of canonical glycolytic metabolites including phosphoenolpyruvate, pyruvate, and fructose-1,6-bisphosphate indicates a disruption of glycolytic flux secondary to mitochondrial perturbation (59). Interestingly, concurrent accumulation of oxidative and non-oxidative PPP intermediates across both evaluated models, notably ribose-5-phosphate, xylulose-5-phosphate, and sedoheptulose-7-phosphate, suggest compensation to respiratory impairment, likely to generate cytosolic NADPH for attempted glutathione recycling, thioredoxin/peroxiredoxin antioxidant systems, and nucleotide synthesis (60,74). The additional accumulation of α-ketoglutarate and glutamic acid in the setting of TCA cycle disruption may further indicating altered glutaminolytic metabolism, potentially redirecting carbons for nucleotide anabolism, lipid synthesis, epigenetic modifications, and NADPH generation for redox homeostasis (59,61). Furthermore, targeting mitochondrial function has particular relevance in ACC as it may address challenging clinical features such as excessive steroid hormone secretion typically resulting in concomitant hypercortisolism, hyperaldosteronism, hyperandrogenism, or hyperestrogenism (2–4,58). These physiological developments often manifest as a wide range of endocrine disorders including Cushing’s syndrome, Conn syndrome, virilization, diabetes mellitus, osteoporosis, and muscle atrophy, collectively contributing to patient comorbidities.(2–4,58) By concurrently targeting tumor metabolism and steroidogenic excess, TR-107 represents a particularly attractive therapeutic approach that uniquely extends beyond conventional cytotoxic strategies.

Ferroptosis is an iron-dependent cell death driven by unrestrained lipid peroxidation and oxidative stress, processes that are closely linked to tumor metabolic state (64,65,67). These intrinsic vulnerabilities have positioned ferroptotic death mechanisms as attractive modalities particularly for tumors with heightened metabolic demand and resistance to traditional therapies (64–67). Our findings extend the mechanistic framework of previous studies and demonstrate that promotion of ferroptosis underlies a central component of TR-107-mediated cytotoxicity in ACC (11). We observed increased intracellular ROS and lipid peroxidation together with substantial remodeling of key regulators governing ferroptotic susceptibility (“the ferroptotic rheostat”), supporting a paradigm in which cellular homeostasis is shifted toward a ferroptosis-permissive state. Specifically, we observe downregulated expression of GPX4, a master mediator of antioxidant defense that maintains redox homeostasis through detoxification of lipid peroxides, ferritin, a principal intracellular iron-sequestering complex that regulates labile iron availability, superoxide dismutase 1, an antioxidant detoxifying superoxide radicals, and SLC7A11, a subunit of the system Xc^-^ cysteine/glutamate antiporter that mediates extracellular cysteine uptake in exchange for intracellular glutamate and thereby supplies the essential substrate for glutathione biosynthesis. Conversely, we observed upregulation of ACSL4, which facilitates ferroptotic sensitivity by enriching membranes with peroxidation-sensitive phospholipids such as phosphatidylethanolamines (11,62,63). The multifactorial ability of TR- 107 to compromise redox homeostasis and induce ferroptosis positions this agent within a new landscape of next-generation precision therapeutics. Identification that TR-107 is not a substrate of the ABCB1 or ABCG2 drug efflux transporters, notorious mediators of multidrug therapeutic resistance, provides further insight into the pharmacological properties of TR-107 while highlighting a potential clinical advantage over conventional anticancer agents in ABCB1/ABCG2-high adrenocortical tumors. This finding is particularly relevant given that the normal adrenal cortex constitutively expresses high levels of ABCB1, an expression pattern that is retained in ACC and further amplified in tumors harboring mutant LOF *p53*, thereby contributing to the intrinsic chemoresistance characteristic of this malignancy (75,76).

Aberrant autocrine and paracrine expression of peptide hormone IGF-2 is a classical hallmark of ACC, occurring in nearly 90% of analyzed adrenocortical tumors (77–79). Additional analyses have revealed IGF-2 mRNA and protein expression to be 10-20 and 8-80 fold higher, respectively, in ACC tissues compared with normal adrenal gland or benign adrenocortical adenomas (77–79). Consequently, integrative pan-genomic expression analyses have identified *IGF2* overexpression as a defining molecular predictor of malignant adrenocortical tumors, with accumulating evidence demonstrating that IGF-2 signaling supports tumor cell proliferation and is associated with poorer clinical outcomes, including shorter progression-free survival (77–80). More recently, pan-cancer genomic analyses of TCGA tumor cohorts have revealed recurrent IGF pathway gene aberrations in ACC, including *IGF1R* overexpression and focal amplification events, thus providing additional evidence of IGF-1R pathway dysregulation in ACC tumorigenesis (81). Given the pervasive dysregulation of the IGF-2/IGF-1R signaling axis in ACC, we investigated whether concomitant targeting of mitochondrial bioenergetics and IGF-1R signaling could enhance therapeutic efficacy. We uncovered robust synergistic antitumor activity between TR-107 and IGF-1R blockade, supporting a combinatory strategy that integrates mitochondrial bioenergetic disruption with targeted inhibition of principal proliferative and survival pathways driving ACC growth and propagation. This approach concurrently exploits distinct yet complementary vulnerabilities and may also overcome compensatory adaptive mechanisms that limit the efficacy of either therapeutic strategy alone. Investigation across ACC xenograft models are currently underway to establish the safety, tolerability, and *in vivo* antitumor pharmacologic profile of TR-107. In summary, these findings establish a strong preclinical rationale for the development of TR-107 as a novel therapeutic agent, either as a monotherapy or as a component of rational combinatory approaches for the treatment of ACC.

## Funding

This work was supported using federal funds from the Center for Cancer Research, National Cancer Institute, National Institutes of Health. The contributions of all NIH authors are considered works of the United States Government. The findings and conclusions presented in this paper are those of the authors and do not necessarily reflect the views of the National Institutes of Health or the U.S. Department of Health and Human Services, nor does mention of trade names, commercial products, or organizations imply endorsement by the U.S. Government.

## Authors’ Disclosures

Edwin J. Iwanowicz has an ownership interest in Madera Therapeutics. All other authors have no competing interests to declare.

## Authors’ Contributions

Conceptualization: G.I.K., S.M.K., M.B., and J.D.R. conceived and designed the study, developed experimental approaches, and established experimental methodologies. Investigation: G.I.K, S.M.K., Y.S.K., H.F., S.N., F.E., and L.L. performed experiments. Analysis: G.I.K., S.M.K., M.B., and J.D.R analyzed and interpreted experimental data. F.E. and A.D. computationally analyzed and interpreted metabolomic and transcriptional datasets. Clinical resources: J.M.H., C.D.H., and J.D.R. provided clinical tumor tissue samples. Resources: E.J.I. and L.M.G. provided the investigational TR-107 compound. Writing: G.I.K. wrote the manuscript. Critical review: S.M.K., M.B., M.I.A., U.W., and J.D.R. critically reviewed and edited the manuscript. Supervision: J.D.R. supervised the study. All authors read and agreed with the final version of the manuscript.

## Acknowledgements

We gratefully acknowledge the philanthropic support of the Adrenal Cancer Collective for the Rare Endocrine and Neuroendocrine Cancer Program at the National Cancer Institute, National Institutes of Health. Additionally, we would like to extend our sincere gratitude to Haojian Li for providing aliquots of liproxstatin-1 and ferrostatin-1, and Takashi Furusawa for providing replication-incompetent HEK293FT-derived retrovirus encoding mito-roGFP2-Orp1. We would like to gratefully acknowledge Nitin Roper for providing mACC3 cells and reagents. Moreover, we would like to gratefully acknowledge Michael Kruhlak, Langston Lim, and Ross Lake at the National Cancer Institute for their invaluable assistance with confocal microscopy. Finally, we would like to gratefully acknowledge all the patients who participated by donating specimens for this study.

## References

1. Fassnacht M, Puglisi S, Kimpel O, Terzolo M. Adrenocortical carcinoma: a practical guide for clinicians. Lancet Diabetes Endocrinol 2025;13(5):438–52 doi 10.1016/S2213-8587(24)00378-4.

2. Shariq OA, McKenzie TJ. Adrenocortical carcinoma: current state of the art, ongoing controversies, and future directions in diagnosis and treatment. Ther Adv Chronic Dis 2021;12:20406223211033103 doi 10.1177/20406223211033103.

3. Libe R, Huillard O. Adrenocortical carcinoma: Diagnosis, prognostic classification and treatment of localized and advanced disease. Cancer Treat Res Commun 2023;37:100759 doi 10.1016/j.ctarc.2023.100759.

4. Chukkalore D, MacDougall K, Master V, Bilen MA, Nazha B. Adrenocortical Carcinomas: Molecular Pathogenesis, Treatment Options, and Emerging Immunotherapy and Targeted Therapy Approaches. Oncologist 2024;29(9):738–46 doi 10.1093/oncolo/oyae029.

5. Tai Y, Shang J. Wnt/beta-catenin signaling pathway in the tumor progression of adrenocortical carcinoma. Front Endocrinol (Lausanne) 2023;14:1260701 doi 10.3389/fendo.2023.1260701.

6. Wang Q, Sun N, Meixner R, Le Gleut R, Kunzke T, Feuchtinger A, et al. Metabolic heterogeneity in adrenocortical carcinoma impacts patient outcomes. JCI Insight 2023;8(16) doi 10.1172/jci.insight.167007.

7. Dedhia PH, Sivakumar H, Rodriguez MA, Nairon KG, Zent JM, Zheng X, et al. A 3D adrenocortical carcinoma tumor platform for preclinical modeling of drug response and matrix metalloproteinase activity. Sci Rep 2023;13(1):15508 doi 10.1038/s41598-023-42659-0.

8. Rios Medrano MA, Bigi MM, Martinez Ponce P, Podesta EJ, Orlando UD. Exposure to anticancer drugs modulates the expression of ACSL4 and ABCG2 proteins in adrenocortical carcinoma cells. Heliyon 2023;9(10):e20769 doi 10.1016/j.heliyon.2023.e20769.

9. Theile D, Haefeli WE, Weiss J. Effects of adrenolytic mitotane on drug elimination pathways assessed in vitro. Endocrine 2015;49(3):842–53 doi 10.1007/s12020-014-0517-2.

10. Ashton TM, McKenna WG, Kunz-Schughart LA, Higgins GS. Oxidative Phosphorylation as an Emerging Target in Cancer Therapy. Clin Cancer Res 2018;24(11):2482–90 doi 10.1158/1078-0432.Ccr-17-3070.

11. Giarrizzo M, LaComb JF, Patel HR, Reddy RG, Haley JD, Graves LM, et al. TR-107, an Agonist of Caseinolytic Peptidase Proteolytic Subunit, Disrupts Mitochondrial Metabolism and Inhibits the Growth of Human Colorectal Cancer Cells. Mol Cancer Ther 2024;23(12):1761–78 doi 10.1158/1535-7163.MCT-24-0170.

12. Fennell EMJ, Aponte-Collazo LJ, Wynn JD, Drizyte-Miller K, Leung E, Greer YE, et al. Characterization of TR-107, a novel chemical activator of the human mitochondrial protease ClpP. Pharmacol Res Perspect 2022;10(4):e00993 doi 10.1002/prp2.993.

13. Le A, Udupa S, Zhang C. The Metabolic Interplay between Cancer and Other Diseases. Trends Cancer 2019;5(12):809–21 doi 10.1016/j.trecan.2019.10.012.

14. DeBerardinis RJ, Chandel NS. Fundamentals of cancer metabolism. Sci Adv 2016;2(5):e1600200 doi 10.1126/sciadv.1600200.

15. Jin P, Jiang JW, Zhou L, Huang Z, Nice EC, Huang CH, et al. Mitochondrial adaptation in cancer drug resistance: prevalence, mechanisms, and management. J Hematol Oncol 2022;15(1) doi ARTN 97 10.1186/s13045-022-01313-4.

16. Martinez-Outschoorn UE, Peiris-Pages M, Pestell RG, Sotgia F, Lisanti MP. Cancer metabolism: a therapeutic perspective. Nat Rev Clin Oncol 2017;14(1):11–31 doi 10.1038/nrclinonc.2016.60.

17. Weyandt JD, Thompson CB, Giaccia AJ, Rathmell WK. Metabolic Alterations in Cancer and Their Potential as Therapeutic Targets. Am Soc Clin Oncol Educ Book 2017;37:825–32 doi 10.1200/EDBK_175561.

18. Fiorillo M, Ozsvari B, Sotgia F, Lisanti MP. High ATP Production Fuels Cancer Drug Resistance and Metastasis: Implications for Mitochondrial ATP Depletion Therapy. Front Oncol 2021;11:740720 doi 10.3389/fonc.2021.740720.

19. Reznik E, Luna A, Aksoy BA, Liu EM, La K, Ostrovnaya I, et al. A Landscape of Metabolic Variation across Tumor Types. Cell Syst 2018;6(3):301–13 e3 doi 10.1016/j.cels.2017.12.014.

20. Bao X, Hou B, Guo Z, Song L, Chen H, Zheng Q, et al. Absolute dynamic and relative static: the relationship of glycolysis and OXPHOS in cancer development. Cell Death Discov 2026;12(1) doi 10.1038/s41420-026-02992-5.

21. Valle S, Alcala S, Martin-Hijano L, Cabezas-Sainz P, Navarro D, Munoz ER, et al. Exploiting oxidative phosphorylation to promote the stem and immunoevasive properties of pancreatic cancer stem cells. Nat Commun 2020;11(1):5265 doi 10.1038/s41467-020-18954-z.

22. Wedam R, Greer YE, Wisniewski DJ, Weltz S, Kundu M, Voeller D, et al. Targeting Mitochondria with ClpP Agonists as a Novel Therapeutic Opportunity in Breast Cancer. Cancers (Basel) 2023;15(7) doi 10.3390/cancers15071936.

23. Kawada M, Inoue H, Ohba S, Hatano M, Amemiya M, Hayashi C, et al. Intervenolin, a new antitumor compound with anti-Helicobacter pylori activity, from Nocardia sp. ML96-86F2. J Antibiot (Tokyo) 2013;66(9):543–8 doi 10.1038/ja.2013.42.

24. Yoshida J, Ohishi T, Abe H, Ohba SI, Inoue H, Usami I, et al. Mitochondrial complex I inhibitors suppress tumor growth through concomitant acidification of the intra- and extracellular environment. iScience 2021;24(12):103497 doi 10.1016/j.isci.2021.103497.

25. Al Assi A, Posty S, Lamarche F, Chebel A, Guitton J, Cottet-Rousselle C, et al. A novel inhibitor of the mitochondrial respiratory complex I with uncoupling properties exerts potent antitumor activity. Cell Death Dis 2024;15(5):311 doi 10.1038/s41419-024-06668-9.

26. Cadassou O, Jordheim LP. OXPHOS inhibitors, metabolism and targeted therapies in cancer. Biochem Pharmacol 2023;211:115531 doi 10.1016/j.bcp.2023.115531.

27. Xu Y, Xue D, Bankhead A, 3rd, Neamati N. Why All the Fuss about Oxidative Phosphorylation (OXPHOS)? J Med Chem 2020;63(23):14276–307 doi 10.1021/acs.jmedchem.0c01013.

28. Beerkens APM, Heskamp S, Reinema FV, Adema GJ, Span PN, Bussink J. Mitochondria Targeting of Oxidative Phosphorylation Inhibitors to Alleviate Hypoxia and Enhance Anticancer Treatment Efficacy. Clin Cancer Res 2025;31(7):1186–93 doi 10.1158/1078-0432.CCR-24-3296.

29. Janku F, LoRusso P, Mansfield AS, Nanda R, Spira A, Wang T, et al. First-in-human evaluation of the novel mitochondrial complex I inhibitor ASP4132 for treatment of cancer. Invest New Drugs 2021;39(5):1348–56 doi 10.1007/s10637-021-01112-7.

30. Nouri K, Feng Y, Schimmer AD. Mitochondrial ClpP serine protease-biological function and emerging target for cancer therapy. Cell Death Dis 2020;11(10):841 doi 10.1038/s41419-020-03062-z.

31. Goncalves MM, Uday AB, Forrester TJB, Currie SQW, Kim AS, Feng Y, et al. Mechanism of allosteric activation in human mitochondrial ClpP protease. Proc Natl Acad Sci U S A 2025;122(16):e2419881122 doi 10.1073/pnas.2419881122.

32. Sauer RT, Fei X, Bell TA, Baker TA. Structure and function of ClpXP, a AAA+ proteolytic machine powered by probabilistic ATP hydrolysis. Crit Rev Biochem Mol Biol 2022;57(2):188–204 doi 10.1080/10409238.2021.1979461.

33. Bonner ER, Waszak SM, Grotzer MA, Mueller S, Nazarian J. Mechanisms of imipridones in targeting mitochondrial metabolism in cancer cells. Neuro Oncol 2021;23(4):542–56 doi 10.1093/neuonc/noaa283.

34. Mabanglo MF, Leung E, Vahidi S, Seraphim TV, Eger BT, Bryson S, et al. ClpP protease activation results from the reorganization of the electrostatic interaction networks at the entrance pores. Commun Biol 2019;2:410 doi 10.1038/s42003-019-0656-3.

35. Baker TA, Sauer RT. ClpXP, an ATP-powered unfolding and protein-degradation machine. Biochim Biophys Acta 2012;1823(1):15–28 doi 10.1016/j.bbamcr.2011.06.007.

36. Cole A, Wang Z, Coyaud E, Voisin V, Gronda M, Jitkova Y, et al. Inhibition of the Mitochondrial Protease ClpP as a Therapeutic Strategy for Human Acute Myeloid Leukemia. Cancer Cell 2015;27(6):864–76 doi 10.1016/j.ccell.2015.05.004.

37. Xu Z, Pokushalov D, Kabir M, Lee Y, Chattopadhyay M, Jenkins EC, et al. Targeting the Mitochondrial Protease ClpP for Anticancer Therapy. J Med Chem 2025;68(20):21377–93 doi 10.1021/acs.jmedchem.5c01315.

38. Reinhardt L, Thomy D, Lakemeyer M, Westermann LM, Ortega J, Sieber SA, et al. Antibiotic Acyldepsipeptides Stimulate the Streptomyces Clp-ATPase/ClpP Complex for Accelerated Proteolysis. mBio 2022;13(6):e0141322 doi 10.1128/mbio.01413-22.

39. Allen JE, Kline CL, Prabhu VV, Wagner J, Ishizawa J, Madhukar N, et al. Discovery and clinical introduction of first-in-class imipridone ONC201. Oncotarget 2016;7(45):74380–92 doi 10.18632/oncotarget.11814.

40. Allen JE, Krigsfeld G, Patel L, Mayes PA, Dicker DT, Wu GS, et al. Identification of TRAIL-inducing compounds highlights small molecule ONC201/TIC10 as a unique anti-cancer agent that activates the TRAIL pathway. Mol Cancer 2015;14:99 doi 10.1186/s12943-015-0346-9.

41. Ishizawa J, Zarabi SF, Davis RE, Halgas O, Nii T, Jitkova Y, et al. Mitochondrial ClpP-Mediated Proteolysis Induces Selective Cancer Cell Lethality. Cancer Cell **2019**;35(5):721–37 e9 doi 10.1016/j.ccell.2019.03.014.

42. Greer YE, Hernandez L, Fennell EMJ, Kundu M, Voeller D, Chari R, et al. Mitochondrial Matrix Protease ClpP Agonists Inhibit Cancer Stem Cell Function in Breast Cancer Cells by Disrupting Mitochondrial Homeostasis. Cancer Res Commun 2022;2(10):1144–61 doi 10.1158/2767-9764.CRC-22-0142.

43. Venneti S, Kawakibi AR, Ji S, Waszak SM, Sweha SR, Mota M, et al. Clinical Efficacy of ONC201 in H3K27M-Mutant Diffuse Midline Gliomas Is Driven by Disruption of Integrated Metabolic and Epigenetic Pathways. Cancer Discov 2023;13(11):2370–93 doi 10.1158/2159-8290.CD-23-0131.

44. U.S. Food and Drug Administration. 2025 FDA grants accelerated approval to dordaviprone for diffuse midline glioma. <https://www.fda.gov/drugs/resources-information-approved-drugs/fda-grants-accelerated-approval-dordaviprone-diffuse-midline-glioma>.

45. Blair HA. Dordaviprone: First Approval. Drugs 2026;86(1):101–9 doi 10.1007/s40265-025-02252-3.

46. Arrillaga-Romany I, Lassman A, McGovern SL, Mueller S, Nabors B, van den Bent M, et al. ACTION: a randomized phase 3 study of ONC201 (dordaviprone) in patients with newly diagnosed H3 K27M-mutant diffuse glioma. Neuro Oncol 2024;26(Supplement_2):S173–S81 doi 10.1093/neuonc/noae031.

47. Thang M, Mellows C, Kass LE, Daglish S, Fennell EMJ, Mann BE, et al. Combining the constitutive TRAIL-secreting induced neural stem cell therapy with the novel anti- cancer drug TR-107 in glioblastoma. Mol Ther Oncol 2024;32(3):200834 doi 10.1016/j.omton.2024.200834.

48. Borges KS, Pignatti E, Leng S, Kariyawasam D, Ruiz-Babot G, Ramalho FS, et al. Wnt/beta-catenin activation cooperates with loss of p53 to cause adrenocortical carcinoma in mice. Oncogene 2020;39(30):5282–91 doi 10.1038/s41388-020-1358-5.

49. Arakawa Y, Jo U, Kumar S, Sun NY, Elloumi F, Thomas A, et al. Activity of the Ubiquitin-activating Enzyme Inhibitor TAK-243 in Adrenocortical Carcinoma Cell Lines, Patient-derived Organoids, and Murine Xenografts. Cancer Res Commun 2024;4(3):834–48 doi 10.1158/2767-9764.CRC-24-0085.

50. Sun Y, Baechler SA, Zhang X, Kumar S, Factor VM, Arakawa Y, et al. Targeting neddylation sensitizes colorectal cancer to topoisomerase I inhibitors by inactivating the DCAF13-CRL4 ubiquitin ligase complex. Nat Commun 2023;14(1):3762 doi 10.1038/s41467-023-39374-9.

51. Di Veroli GY, Fornari C, Wang D, Mollard S, Bramhall JL, Richards FM, et al. Combenefit: an interactive platform for the analysis and visualization of drug combinations. Bioinformatics (Oxford, England) 2016;32(18):2866–8 doi 10.1093/bioinformatics/btw230.

52. Schneider CA, Rasband WS, Eliceiri KW. NIH Image to ImageJ: 25 years of image analysis. Nat Methods 2012;9(7):671–5 doi 10.1038/nmeth.2089.

53. Geissmann Q. OpenCFU, a new free and open-source software to count cell colonies and other circular objects. PLoS One 2013;8(2):e54072 doi 10.1371/journal.pone.0054072.

54. Hanson GT, Aggeler R, Oglesbee D, Cannon M, Capaldi RA, Tsien RY, et al. Investigating mitochondrial redox potential with redox-sensitive green fluorescent protein indicators. J Biol Chem 2004;279(13):13044–53 doi 10.1074/jbc.M312846200.

55. Teixeira RB, Karbasiafshar C, Sabra M, Abid MR. Optimization of mito-roGFP protocol to measure mitochondrial oxidative status in human coronary artery endothelial cells. STAR Protoc 2021;2(3):100753 doi 10.1016/j.xpro.2021.100753.

56. Robey RW, Lin B, Qiu J, Chan LL, Bates SE. Rapid detection of ABC transporter interaction: potential utility in pharmacology. J Pharmacol Toxicol Methods 2011;63(3):217–22 doi 10.1016/j.vascn.2010.11.003.

57. Mizdrak M, Ticinovic Kurir T, Bozic J. The Role of Biomarkers in Adrenocortical Carcinoma: A Review of Current Evidence and Future Perspectives. Biomedicines 2021;9(2) doi 10.3390/biomedicines9020174.

58. Ghosh C, Hu J, Kebebew E. Advances in translational research of the rare cancer type adrenocortical carcinoma. Nat Rev Cancer 2023;23(12):805–24 doi 10.1038/s41568-023-00623-0.

59. Pavlova NN, Zhu J, Thompson CB. The hallmarks of cancer metabolism: Still emerging. Cell Metab 2022;34(3):355–77 doi 10.1016/j.cmet.2022.01.007.

60. TeSlaa T, Ralser M, Fan J, Rabinowitz JD. The pentose phosphate pathway in health and disease. Nat Metab 2023;5(8):1275–89 doi 10.1038/s42255-023-00863-2.

61. Yoo HC, Yu YC, Sung Y, Han JM. Glutamine reliance in cell metabolism. Exp Mol Med 2020;52(9):1496–516 doi 10.1038/s12276-020-00504-8.

62. Tang D, Chen X, Kang R, Kroemer G. Ferroptosis: molecular mechanisms and health implications. Cell Res 2021;31(2):107–25 doi 10.1038/s41422-020-00441-1.

63. Zhang XD, Liu ZY, Wang MS, Guo YX, Wang XK, Luo K, et al. Mechanisms and regulations of ferroptosis. Front Immunol 2023;14:1269451 doi 10.3389/fimmu.2023.1269451.

64. Li J, Jia YC, Ding YX, Bai J, Cao F, Li F. The crosstalk between ferroptosis and mitochondrial dynamic regulatory networks. Int J Biol Sci 2023;19(9):2756–71 doi 10.7150/ijbs.83348.

65. Liu Y, Lu S, Wu LL, Yang L, Yang L, Wang J. The diversified role of mitochondria in ferroptosis in cancer. Cell Death Dis 2023;14(8):519 doi 10.1038/s41419-023-06045-y.

66. Zhou Q, Meng Y, Li D, Yao L, Le J, Liu Y, et al. Ferroptosis in cancer: From molecular mechanisms to therapeutic strategies. Signal Transduct Target Ther 2024;9(1):55 doi 10.1038/s41392-024-01769-5.

67. Wang T, Zhou X, Yin X, Zhang A, Fan Y, Chen K, et al. From mitochondrial dysregulation to ferroptosis: Exploring new strategies and challenges in radioimmunotherapy (Review). Int J Oncol 2025;67(3) doi 10.3892/ijo.2025.5781.

68. Lee D, Jeong HS, Hwang SY, Lee YG, Kang YJ. ABCB1 confers resistance to carboplatin by accumulating stem-like cells in the G2/M phase of the cell cycle in p53(null) ovarian cancer. Cell Death Discov 2025;11(1):132 doi 10.1038/s41420-025-02435-7.

69. Engle K, Kumar G. Cancer multidrug-resistance reversal by ABCB1 inhibition: A recent update. Eur J Med Chem 2022;239:114542 doi 10.1016/j.ejmech.2022.114542.

70. Bahar ME, Kim HJ, Kim DR. Targeting the RAS/RAF/MAPK pathway for cancer therapy: from mechanism to clinical studies. Signal Transduct Target Ther 2023;8(1):455 doi 10.1038/s41392-023-01705-z.

71. Glaviano A, Foo ASC, Lam HY, Yap KCH, Jacot W, Jones RH, et al. PI3K/AKT/mTOR signaling transduction pathway and targeted therapies in cancer. Mol Cancer 2023;22(1):138 doi 10.1186/s12943-023-01827-6.

72. Miller WL, Auchus RJ. The molecular biology, biochemistry, and physiology of human steroidogenesis and its disorders. Endocr Rev 2011;32(1):81–151 doi 10.1210/er.2010-0013.

73. Scicluna P, Caramuta S, Kjellin H, Xu C, Frobom R, Akhtar M, et al. Altered expression of the IGF2-H19 locus and mitochondrial respiratory complexes in adrenocortical carcinoma. Int J Oncol 2022;61(5) doi 10.3892/ijo.2022.5430.

74. Stancill JS, Corbett JA. The Role of Thioredoxin/Peroxiredoxin in the beta-Cell Defense Against Oxidative Damage. Front Endocrinol (Lausanne) 2021;12:718235 doi 10.3389/fendo.2021.718235.

75. Lopez JP, Brivio E, Santambrogio A, De Donno C, Kos A, Peters M, et al. Single-cell molecular profiling of all three components of the HPA axis reveals adrenal ABCB1 as a regulator of stress adaptation. Sci Adv 2021;7(5) doi 10.1126/sciadv.abe4497.

76. Skinner KT, Palkar AM, Hong AL. Genetics of ABCB1 in Cancer. Cancers (Basel) 2023;15(17) doi 10.3390/cancers15174236.

77. Drelon C, Berthon A, Val P. Adrenocortical cancer and IGF2: is the game over or our experimental models limited? J Clin Endocrinol Metab 2013;98(2):505–7 doi 10.1210/jc.2012-3310.

78. Pereira SS, Monteiro MP, Costa MM, Moreira A, Alves MG, Oliveira PF, et al. IGF2 role in adrenocortical carcinoma biology. Endocrine 2019;66(2):326–37 doi 10.1007/s12020-019-02033-5.

79. Guillaud-Bataille M, Ragazzon B, de Reynies A, Chevalier C, Francillard I, Barreau O, et al. IGF2 promotes growth of adrenocortical carcinoma cells, but its overexpression does not modify phenotypic and molecular features of adrenocortical carcinoma. PLoS One 2014;9(8):e103744 doi 10.1371/journal.pone.0103744.

80. de Reynies A, Assie G, Rickman DS, Tissier F, Groussin L, Rene-Corail F, et al. Gene expression profiling reveals a new classification of adrenocortical tumors and identifies molecular predictors of malignancy and survival. J Clin Oncol 2009;27(7):1108–15 doi 10.1200/JCO.2008.18.5678.

81. Wang P, Mak VC, Cheung LW. Drugging IGF-1R in cancer: New insights and emerging opportunities. Genes Dis 2023;10(1):199–211 doi 10.1016/j.gendis.2022.03.002.

