## Supplementary Figures for "TR-107, a novel mitochondrial ClpP agonist, induces robust antitumor activity against preclinical models of adrenocortical carcinoma"

### Supplementary Figure S1

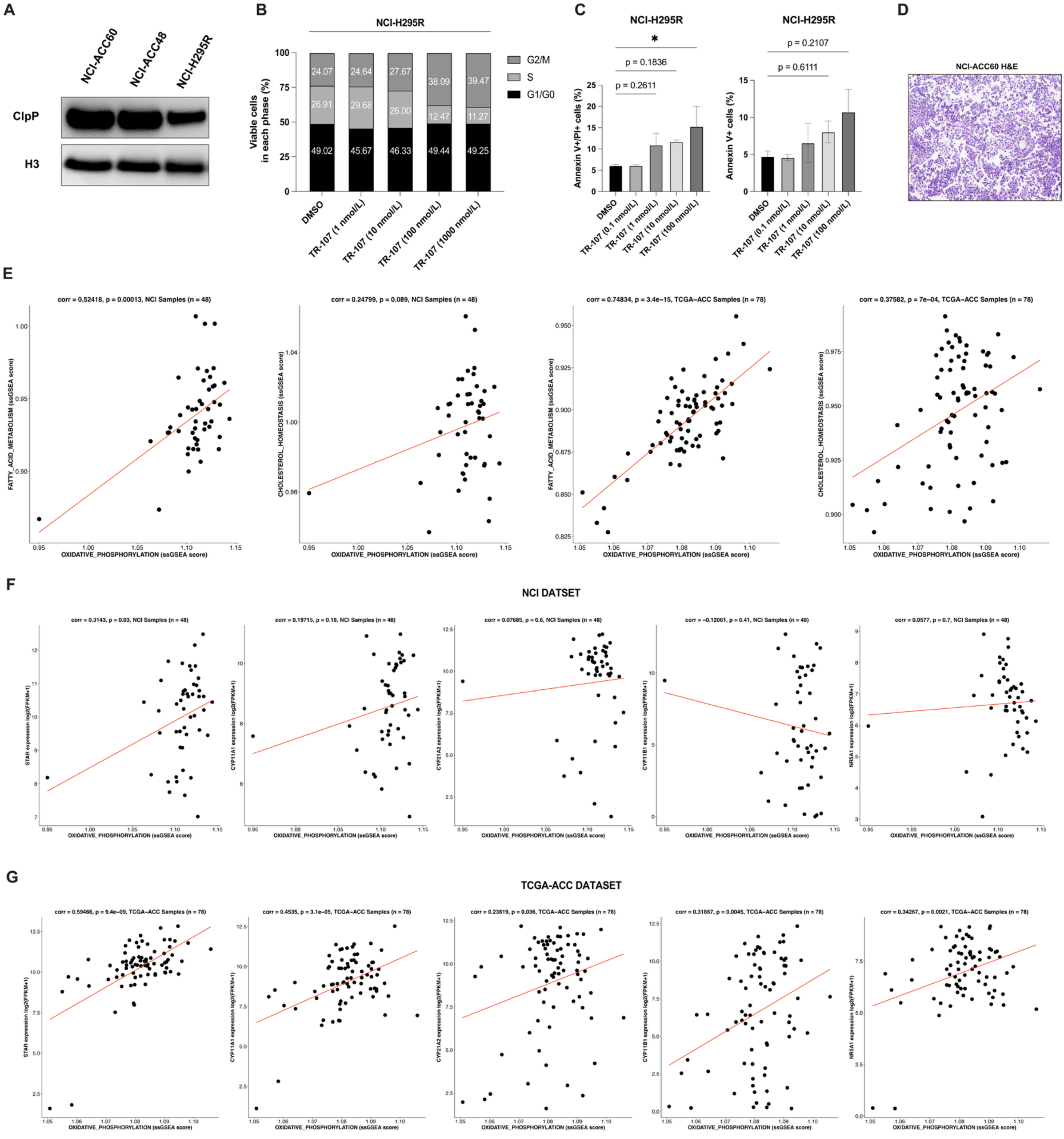

**A,** Immunoblot analysis of ClpP expression in representative ACC models NCI-H295R, NCI-ACC48 PDOs, and NCI-ACC60 PDOs. **B,** Fractions of viable NCI-H295R cells in each cell cycle phase following 48-hour TR-107 treatment. **C,** Quantification of Annexin V-positive/propidium iodide-positive and Annexin V-positive NCI-H295R cells following 48-hour TR-107 treatment. **D,** Hematoxylin & eosin (H&E) staining of NCI-ACC60 PDOs. **E,** Pearson correlation analyses between fatty acid metabolism, cholesterol homeostasis ssGSEA enrichment scores. **F** and **G,** the mRNA expression of key steroidogenic proteins relative to oxidative phosphorylation ssGSEA enrichment scores in NCI-ACC (**F**) and TCGA-ACC (**G**) tumor specimens. n.s., not significant; *, $P < 0.05$.

#### Supplementary Figure S2.

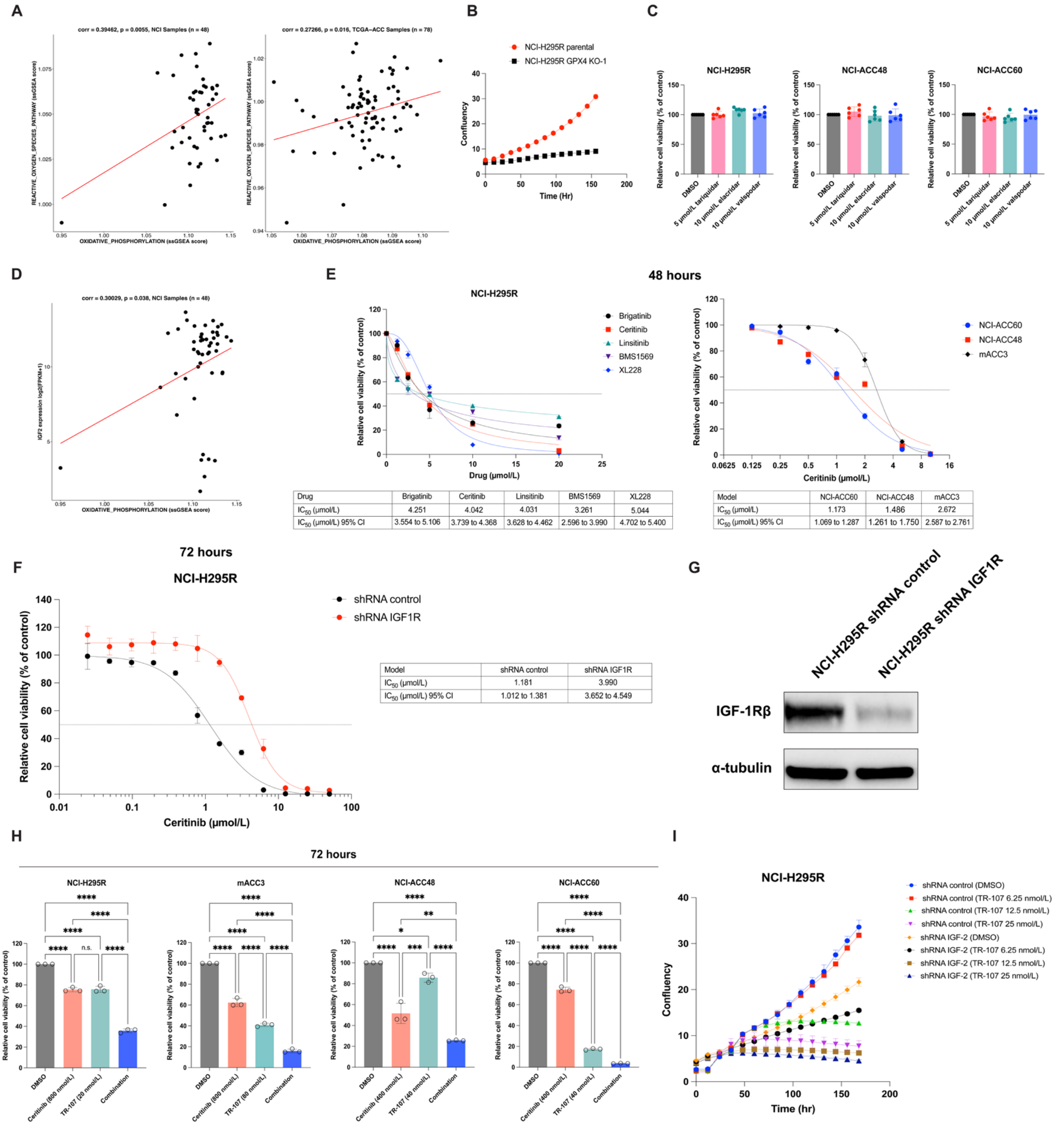


| Drug | Brigatinib | Ceritinib | Linsitinib | BMS1569 | XL228 |
|---|---|---|---|---|---|
| $IC_{50}$ (μmol/L) | 4.251 | 4.042 | 4.031 | 3.261 | 5.044 |
| $IC_{50}$ (μmol/L) 95% CI | 3.554 to 5.106 | 3.739 to 4.368 | 3.628 to 4.462 | 2.596 to 3.990 | 4.702 to 5.400 |

| Model | NCI-ACC60 | NCI-ACC48 | mACC3 |
|---|---|---|---|
| $IC_{50}$ (μmol/L) | 1.173 | 1.486 | 2.672 |
| $IC_{50}$ (μmol/L) 95% CI | 1.069 to 1.287 | 1.261 to 1.750 | 2.587 to 2.761 |

| Model | shRNA control | shRNA IGF1R |
|---|---|---|
| $IC_{50}$ (μmol/L) | 1.181 | 3.990 |
| $IC_{50}$ (μmol/L) 95% CI | 1.012 to 1.381 | 3.652 to 4.549 |

**A,** Pearson correlation analysis between reactive oxygen species (ROS) ssGSEA enrichment scores and oxidative phopshorylation ssGSEA enrichment score in NCI-ACC and TCGA-ACC tumor specimens. **B,** Longitudinal basal cell proliferation of NCI-H295R GPX4 KO cells compared to NCI-H295R parental cells. Data reported as mean ± SEM, $n = 3$ technical replicates. **C,** Relative cell viability of NCI-H295R cells cultured with ABC transporter inhibitors tariquidar (5 µmol/L), elacridar (10 µmol/L), and valspodar (10 µmol/L) alone. Data reported as mean ± SD, $n = 6$ technical replicates. **D,** Pearson correlation analysis between *IGF2* expression and oxidative phopshorylation ssGSEA enrichment score in NCI-ACC and TCGA-ACC tumor specimens. **E,** Relative cytotoxic effects of variable IGF-1R inhibitors following 48-hour treatment in NCI-H295R, mACC3, NCI-ACC48, and NCI-ACC60 models. Data reported as mean ± SD, $n = 3$ technical replicates. **F,** Relative cytotoxic effects of 72-hour ceritinib treatment on NCI-H295R *IGF1R* shRNA and control shRNA cells. **G,** Immunoblot validation of NCI-H295R *IGF1R* shRNA and control shRNA cells. **H,** Relative cytotoxic effects of TR-107 plus ceritinib combinatory treatment in NCI-H295R, mACC3, NCI-ACC48, and NCI-ACC60 models following 72-hour treatment. Data reported as mean ± SD, $n = 3$ technical replicates. **I,** Longitudinal cell proliferation of NCI-H295R *IGF2* shRNA and NCI-H295R control shRNA cells following treatment with variable concentrations. Data reported as mean ± SEM, $n = 3$ technical replicates. n.s., not significant; *, $P < 0.05$; **, $P < 0.01$; ***, $P < 0.001$; ****, $P < 0.0001$.
